# Application of computational data modeling to a large-scale population cohort assists the discovery of specific nutrients that influence beneficial human gut bacteria *Faecalibacterium prausnitzii*

**DOI:** 10.1101/2022.12.19.518690

**Authors:** Shaillay Kumar Dogra, Adrien Dardinier, Fabio Mainardi, Léa Siegwald, Simona Bartova, Caroline le Roy, Chieh Jason Chou

## Abstract

*Faecalibacterium prausnitzii* (*F. prausnitzii*) is a bacterial taxon of the human gut with anti-inflammatory properties and negative associations with chronic inflammatory conditions. *F. prausnitzii* may be one of key species contributing to the effects of healthy eating habits, and yet little is known about the nutrients that enhance the growth of *F. prausnitzii* other than simple sugars and fibers. Here we combined dietary and microbiome data from the American Gut Project (AGP) to identify nutrients that may be linked to the relative abundance of *F. prausnitzii*. Using a machine learning approach in combination with univariate analyses, we identified that sugar alcohols, carbocyclic sugar and vitamins may contribute to *F. prausnitzii* growth. We next explored the effects of these nutrients on the growth of two *F. prausnitzii* strains *in vitro* and observed strain dependent growth patterns on the nutrient tested. In the context of a complex community using *in vitro* fermentation, none of the tested nutrients and nutrient combinations exerted a significant growth-promoting effect on *F. prausnitzii* due to high variability in batch responses. A positive association between *F. prausnitzii* and butyrate concentrations was observed. Future nutritional studies aiming to increase relative abundance of *F. prausnitzii* should explore a personalized approach accounting for strain-level genetic variations and community-level microbiome composition.

## Introduction

*Faecalibacterium prausnitzii* (*F. prausnitzii*) belongs to the *Ruminococcaceae* family (phylum Firmicutes) and is one of most abundant bacteria in the human gut [1]. It has been demonstrated to associate with (severity or incidence) of different diseases in humans and to play a causative role in mouse models [2]. Reduced abundance of *F. prausnitzii* has consistently been found in disease conditions such as inflammatory bowel disease (IBD) [3], irritable bowel syndrome (IBS), metabolic syndrome and diabetes [4-7], NAFLD and NASH [8], colorectal cancer (CRC) [9], obesity and frailty [10]. Functionally, *F. prausnitzii* contributes to the modulation of the immune system and attenuation of inflammation through multiple mechanisms that can work independently or synergistically. More precisely, butyrate produced by *F. prausnitzii* and other butyrate producing bacteria reduces intestinal mucosal inflammation by inhibiting NF-κβ activation, upregulating peroxisome proliferator-activated receptor-γ expression and inhibiting interferon-γ expression [10]. In addition, *F. prausnitzii* modulates inflammatory signals by releasing immune-suppressing molecules such as salicyclic acid [11] and microbial anti-inflammatory molecules (MAM) [12]. The therapeutic potential of *F. prausnitzii* through secretion of microbial anti-inflammatory molecules has been demonstrated in a mouse model of IBD [13]. Together with association-based evidence from observational and clinical studies, scientists have argued for the use of *F. prausnitzii* as a probiotic [10].

According to ISAPP, the definition of probiotics is “live microorganisms that, when administered in adequate amounts, confer a health benefit on the host” [14]. *F. prausnitzii* is currently not accepted as a probiotic due to the lack of clinical evidence on its safety and efficacy. Extreme oxygen sensitivity of *F. prausnitzii* imposes practical challenges to the production, transportation, storage, and manufacturing of probiotic products to be evaluated in a clinical setting. Alternatively, relative abundance of *F. prausnitzii* in the human gut can be affected by multiple factors such as antibiotic usage [15] and diet [16, 17]. More precisely, some food ingredients have been shown to increase the abundance of *F. prausnitzii* in humans. Thus, a prebiotic approach aiming to enhance health through increasing the abundance of commensal *F. prausnitzii* could be a viable strategy.

Indeed, *F. prausnitzii* relative abundance in the human gut appears to be associated to diet healthiness (based on a healthy eating index) [18]. More specifically, consumption of prebiotic-type ingredients such as inulin and fructo-oligosaccharides were found to increase *F. prausnitzii* in obese women [19], IBS patients [20] and healthy individuals [21]. Treatment with polydextrose and chickpea oligosaccharides (raffinose) also lead to the increase of *F. prausnitzii* abundance in fecal communities of healthy subjects [22, 23]. Yet, deconvoluting the effects of individual nutrients or food items on *F. prausnitzii* in the gut from the rest of the diet remains challenging.

Thus, our aim was to identify nutrients that could be used to boost *F. prausnitzii* abundance in the human gut. To this end, we applied a machine learning algorithm on dietary records and 16S rRNA gene sequencing data collected on 3816 participants of the American Gut Project (AGP) to identify new nutrients that may link to the relative abundance of *F. prausnitzii*. We next evaluated the impact of selected nutrients on the growth of *F. prausnitzii in vitro* using pure culture of single strains and fermentation of human fecal communities.

## Materials and Methods

### Data

The intersection of three datasets [metadata, microbiota and VioScreen food frequency questionnaires (FFQ)] from AGP [24] was used in this study and represented a sample size of n=3816 (Figure S1). Raw 16S rRNA gene sequencing data from stool samples was downloaded on September 4^th^, 2019 from the QIITA repository https://qiita.ucsd.edu/study/description/10317 [25]. Data was processed following the same analytical steps as in the original publication [24]. Briefly, raw sequencing reads were firstly denoised and OTUs were generated using deblur v1.0.2 [26]. Then, OTUs matching bacteria potentially blooming under room temperature storage conditions were removed following the instructions of https://github.com/knightlab-analyses/bloom-analyses. Multiple rarefactions were performed ten times at a threshold of 1250 sequences per sample. Finally, representative sequences of each OTU were annotated using the QIIME2 v. 2017.4 RDP classifier on Greengenes 99% v. 13.8 [27]. Nutrient data as provided through VioScreen FFQ analysis was downloaded on July 31^st^, 2019 from the AGP data FTP site http://ftp.microbio.me/AmericanGut/raw-vioscreen/vioscreen_dump.tsv.gz. Coded names and full description of the nutrients and the corresponding units of nutrients are shown in Table S1. The metadata file “10317_20220801-114642.txt” was downloaded on September 15^th^, 2022 from the QIITA repository https://qiita.ucsd.edu/study/description/10317.

### Modelling to predict the abundance of F. prausnitzii using nutrient intake data

Predictive models were built to determine the relative abundance of *F. prausnitzii* of an individual subject based on nutrient intake values. In particular, the model predicted the *F. prausnitzii* relative abundance by several nutrient feature parameters to determine the *F. prausnitzii* abundance category of the subject as defined in Table S2 (e.g., “Low” or “notLow”; “High” or “notHigh”; “Low” or “High”). A cube-root transformation was applied on the *F. prausnitzii* abundances to make them normally distributed before binning them into these different categories (Figure S2).

Data were split into a training set “Train” (80%) and a testing set “holdout/Test set” (20%). For optimal model performance, we used random downsampling to match the number of subjects between the abundance groups. For example, we randomly downsampled in notLow group to match the subject number of low group. The Train set was used by different machine learning algorithms (RandomForests, XGBoost) to train a model. The learning from the data was done in a cross-validated manner where Train data was split into partitions with some parts used for training the model and other for internal testing (Repeated k-Fold Cross-Validation, i.e., 3-folds, 3-repeats). The holdout/Test set was used only for checking the performance of the final trained model and was not used during the model training phase. A total of 9 models (model A to model I) were made, and the cut offs used to define the groups, the type of machine learning algorithm used and other parameters for each of the model are provided in Table S2.

Receiver Operating Characteristic (ROC) curves were generated for these models and Area Under the Curve (AUC) reported. The best performing model was then selected from the different binning categories of Low vs. notLow, High vs. notHigh, and Low vs. High (Table S2).

### Culture conditions for testing selected nutrients

*F. prausnitzii* strains A2-165 and 27768 were obtained from the Deutsche Sammlung von Mikroorganismen und Zellkulturen GmbH (DSMZ) and American Type Culture Collection (ATCC), respectively. Culture of *F. prausnitzii* followed the method of Duncan et al. [28] using Hungate culture tubes in an anaerobic chamber (H_2_:CO_2_:N_2_, 5:10:85%, Type B, Coy Laboratory Products, Grass Lake, MI, USA). To prepare the working cultures, lyophilized *F. prausnitzii* (ATCC 27768) were enumerated anaerobically with 20 mL ATCC media 2107 consisting in trypose 10 g/L, beef extract 10 g/L, yeast extract 3g/L, dextrose 5 g/L, NaCl 5 g/L, starch 1 g/L, L-cysteine HCl 0.5 g/L, sodium acetate 3 g/L, resazuim (0.025%) 4 mL/L in dd water for 3 days. For each experiment, 0.5 mL of homogenized live liquid culture was added to 9 mL freshly prepared YCFA media in a Hungate tube under an anaerobic condition. Growth of bacteria was evaluated by the measurement of optical density at 600 nm (Biowave WPA CO8000 – WPA Cambridge, UK) after incubation at 37^°^C on a rotating platform inside of an anaerobic chamber.

Glucose (SigmaAldrich) at 10 mM was used as a control carbohydrate source to verify that the strains grew under the assay conditions. To test the ability of *F. prausnitzii* to grow on different carbon sources, glucose was replaced by the same concentration of inositol, sorbitol, erythritol, pinitol or xylitol prepared with sterile ddH_2_O that was pre-flushed with N_2_ gas. The effect of vitamins was tested at the final concentration of 1 ug/10 mL for vitamin B5 and B6, at 0.05 ug/10 mL for vitamin B12 or 0.1 ug/10 mL for vitamin A and D in the YCFA media with either glucose or inositol as the main carbon source. YCFA media were autoclaved at 121°C for 15 min and were transferred to an anaerobic chamber till use. L-cysteine-HCl, thiamine hydrochloride (T4625-5G, SigmaAldrich), and riboflavin (R4500, SigmaAldrich) were first sterile filtered (0.2 um, Media bottle filtration unit with PES membrane, VWR No. 514-0297) and added to the media prior to each experiment and the pH was adjusted to 6.7 with NaOH immediately before the start of the experiment.

### Batch fermentation

Fecal samples of healthy volunteers were collected under a protocol approved by Lausanne ethical committee (CER-VD) (authorization number: 2020-00304). Preparation of stool samples for *in vitro* fermentation followed the procedure described by Van den Abbeele et al. [29]. Freshly collected stool samples were placed in an air-tight jar equipped with AnaeroGen™ to reduce the exposure of ambient oxygen. Once inside of an anaerobic chamber (Coy Laboratory Products, Grass Lake, MI, USA), fecal materials were diluted 10 times (w/v) in anaerobic phosphate buffer (0.1M of NaH_2_PO_4_ and 0.1M of Na_2_HPO_4_ in 2:1 ratio) containing 10% glycerol and the aliquots of fecal stocks (25 mL) were stored at - 80^°^C for later use. *In vitro* fermentation experiment carried in a Hungate tubes where 0.25 mL of fecal stock solution was inoculated to 10 mL of a casitone supplemented oligotrophic medium: casitone (10g/L), L-cysteine (0.05%), NaCl (8g/L), KCl (0.2g/L), Na2HPO4 (1.15g/L), KH2PO4 (0.2g/L) at pH 7.3 as starting of fermentation [30]. Inulin or inositol at 10 mM was added in the basic culture media, and vitamins B5, B6, B12, A and D were included in the relevant groups at the same concentrations as pure culture experiments mentioned above. Samples were collected at time 0, 6h, 24h and 48h from the start of the experiment for the quantification of *F. prausnitzii* and metabolomic analysis.

### Bacterial DNA extraction

Bacterial DNA was extracted using QIAamp Fast-DNA Mini Kit (Qiagen, no: 51604, Germany) following the manufacturer’s recommended procedure. In short, *in vitro* fermentation samples (1 ml) were mixed an equal amount of InhibitEX buffer in a Lysing Matrix B tube before two steps of homogenization with Fastprep (M.P. Biomedicals, Irvine, California, USA). Lysate was further prepared by centrifugation, proteolytic digestion with Protease K and incubation (10 min). Then DNA was extracted and purified with QIAamp spin column. The concentration of resulting DNA was measured by spectrophotometric or fluorescent methods (DropSense96, Qubit 2.0, PicoGreen). DNA samples were stored at -20^°^C before the quantification of *F. prausnitzii*.

### Quantification of total bacteria and F. prausnitzii by real time PCR

Quantification of total bacteria and *F. prausnitzii* was performed with real time PCR using the ABI-PRISM 7700 Sequence Detection System (Applied Biosystems) in duplicates. Quantification of total bacteria was performed in a total volume of 25 µl reagent mix using the Perfecta MasterMix (Quantabio, PerfeCta® qPCR ToughMix® ROX, Beverley MA, USA), containing 300 nM of each of the forward (f: TCCTACGGGAGG CAGCAGT) and reverse primers (r: GGACTACCAGGG TATCTAATCCTGTT) and 175 nM of fluorogenic probe (FAM-CGTATTACCGCG GCTGCTGGCAC-BHQ) as described by Nadkarni et al [31]. The amplifications of DNA were 95°C for 10 min and 50 cycles of 95°C for 15 s and 60°C for 1 min. Detection of *F. prausnitzii* follows the method described by Lopez-Siles et al. [32]. In short, PCR reactions were carried out in 20 µl containing TaqMan Universal PCR Master Mix, 300 nM of each of the forward (Fpra 428 F TGTAAACTCCTGTTGTTGAGGAAGATAA) and reverse (Fpra 583 R GCGCTCCCTTTACACCCA) primers and 200 nM of Probe (Fpra 493 PR 6FAM-CAAGGAAGTGACGGCTAACTACGTGCCAG-TAMRA). Data analysis made use of Sequence Detection Software version 1.6.3 supplied by Applied Biosystems.

### ^1^H-NMR metabolomics

Frozen samples from *in vitro* fermentation were thawed at room temperature before centrifugation for 10 min at 10,000 x g at 4°C. The supernatants (300 *μ*L) were then mixed with 300 *μ*L sodium phosphate buffer 0.075 M at pH 7.4, vortex mixed and 560 *μ*L were transferred to 5 mm NMR tubes. The samples were then analyzed by 1D ^1^H-NMR in a 600 MHz Bruker spectrometer at 300 K. A set of 2D NMR experiments (^1^H J-Resolved, ^1^H-^1^H COSY and ^1^H-^13^C HSQC) were acquired for selected samples to aid metabolite identification. All NMR spectral acquisition and pre-processing were done under the control of TopSpin 4.0.9 (Bruker BioSpin, Rheinstetten, Germany), and the automated submission of a sequence of samples was performed using ICON-NMR 5 (Bruker BioSpin, Rheinstetten, Germany). Metabolite annotation was performed by comparing metabolite signals to those of Bruker BIOREFCODE library from public databased Human Metabolome Database (HMDB) [33].

To analyze the data, 1D NMR spectra were imported into R statistical software environment (version 4.1.1) [34] using the AlpsNMR package [35] and intensities and chemical shifts were interpolated to get a consistent shared ppm axis for all spectra between -0.5 and 10 ppm. Residual signal of water (4.70 to 4.9 ppm) was removed. NMR targeted peak integration was performed using a numeric integration automated routine in R statistical software. The integrated data were log transformed prior statistical analysis. Metabolic profile was visualized by a principal component analysis (PCA) performed using unit-variance scaling.

### General statistical analysis

Comparisons of *F. prausnitzii* relative abundance between groups were performed with Kruskal-Wallis rank sum test followed by post hoc Dunn test. Wilcoxon rank sum test was used to compare the intake of nutrients between low and notlow *F. prausnitzii* categories and Benjamini-Hochberg method was applied to control the false discovery rate (0.05). Descriptive statistics on the Healthy Eating Index-2010 were based on data published by the USDA (https://fns-prod.azureedge.us/sites/default/files/media/file/HEI2010_Age_Groups_2011_2012.pdf). The above analyses were performed using the R statistical software, version 4.1.1.

Data are expressed as mean +/- SEM for data from *in vitro* culture and fermentation experiments. Comparisons between the groups were examined with One-way ANOVA followed by Tukey multiple comparisons test using GraphPad Prism version 9.2.0 for Windows (GraphPad Software, San Diego, California USA, www.graphpad.com).

## Results

### Characteristics of the study subjects

We used microbiome and dietary data collected on 3816 individuals from the AGP cohort [36]. The full description of the cohort is shown in Table S3. There is higher percentage of female (59.4%) than male (38.9%), and participant self-reported country of residence is primarily from the US (43.3%), followed by UK (21.7%) and Australia (1.5%). The average age of the study population is 51.3 ± 15.6 years old (mean ± SD), and a large portion falls into normal BMI category (54.9%) with some being overweight (28.9%), obese (9.5%) and underweight (4.4%). In terms of dietary preference, 76.8% declared as omnivore and remaining subjects follow vegetarian (4.8%), vegan (3.2%) and other e.g., tribal diets (Table S3). Quality of nutrition intake as measured by Health Eating Index (HEI) is 66.34 ± 1.38 for children (2-17 yrs, n=68), 70.8 for adults (18-64 yrs, n=2686), and 71.54 ± 0.32 for older adults (≥65 yrs, n=853). The HEI scores appear to be higher in all age groups of AGP subjects than age matched general US population (NHANES 2011-2012), although a statistical comparison is not possible due to the different methods in collecting dietary intake information (Table 1). Interestingly, we found significant declines of *F. prausnitzii* abundance with age (Kruskal-Wallis rank sum test, p=6.9 × 10^−5^, Dunn test 20s vs. 50s adj p=0.04; 20s vs. 60s adj p=0.03, Figure S3).

**Table 1.**
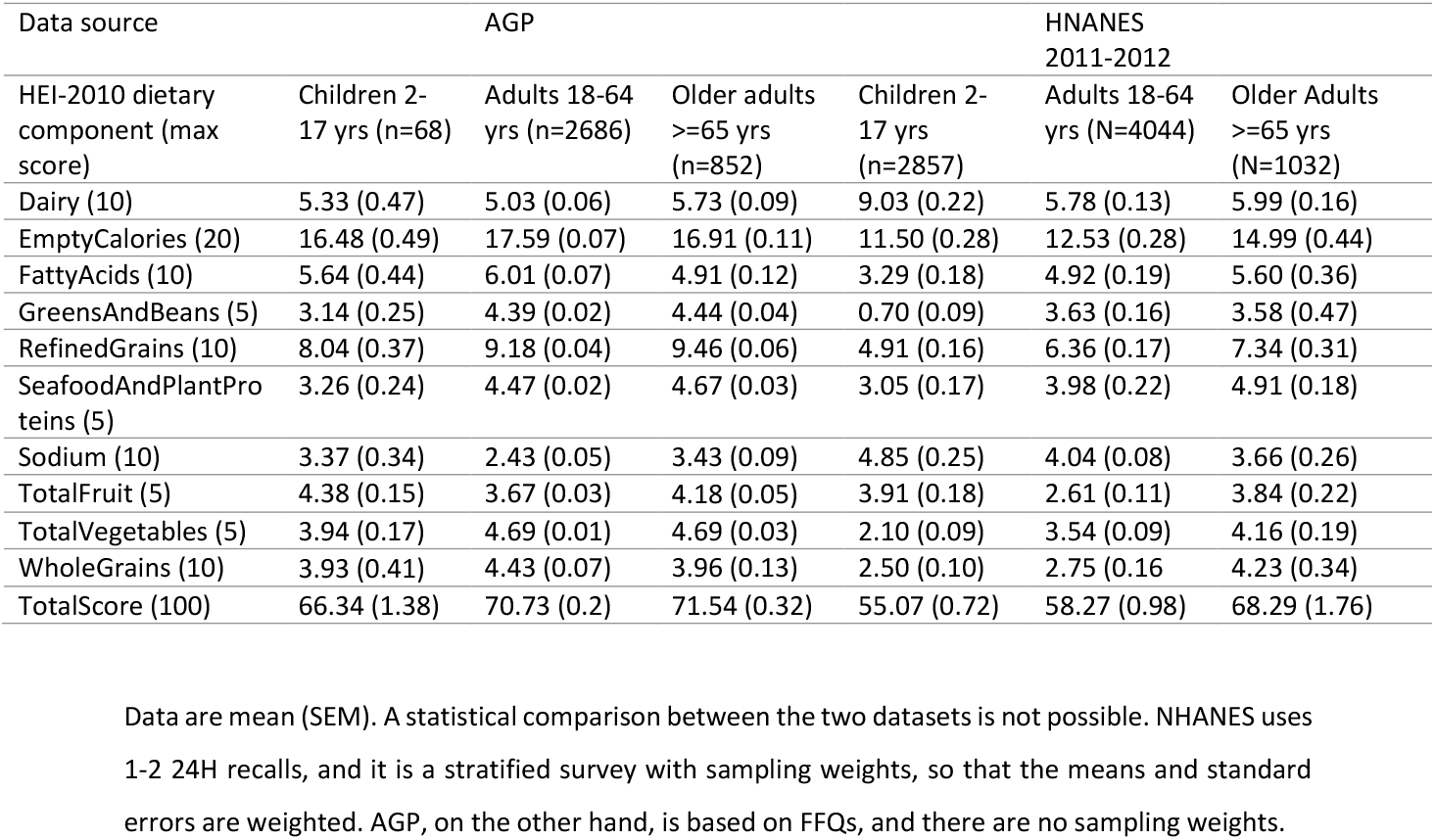
Comparison of diet quality measured as healthy eating index between AGP subjects and general American population (NHANES 2011-2012)

### Discovery of nutrients associated with the abundance of F. prausnitzii

To identify nutrients that can predict the relative abundance of *F. prausnitzii* in the gut ecosystem as estimated from fecal sampling, random forest and XGBoost Machine Learning models with 3-folds cross validation were generated using 251 nutrition-related features extracted from FFQs. Among all considered models, model E (low vs notlow with cut off based on mean – 1SD, Table S2) performed the best with an AUC-ROC of 0.65 ± 0.02 for the training (n= 896) and 0.68 (n=764) for the test set (Figure 1A and 1B).

**Figure 1.**
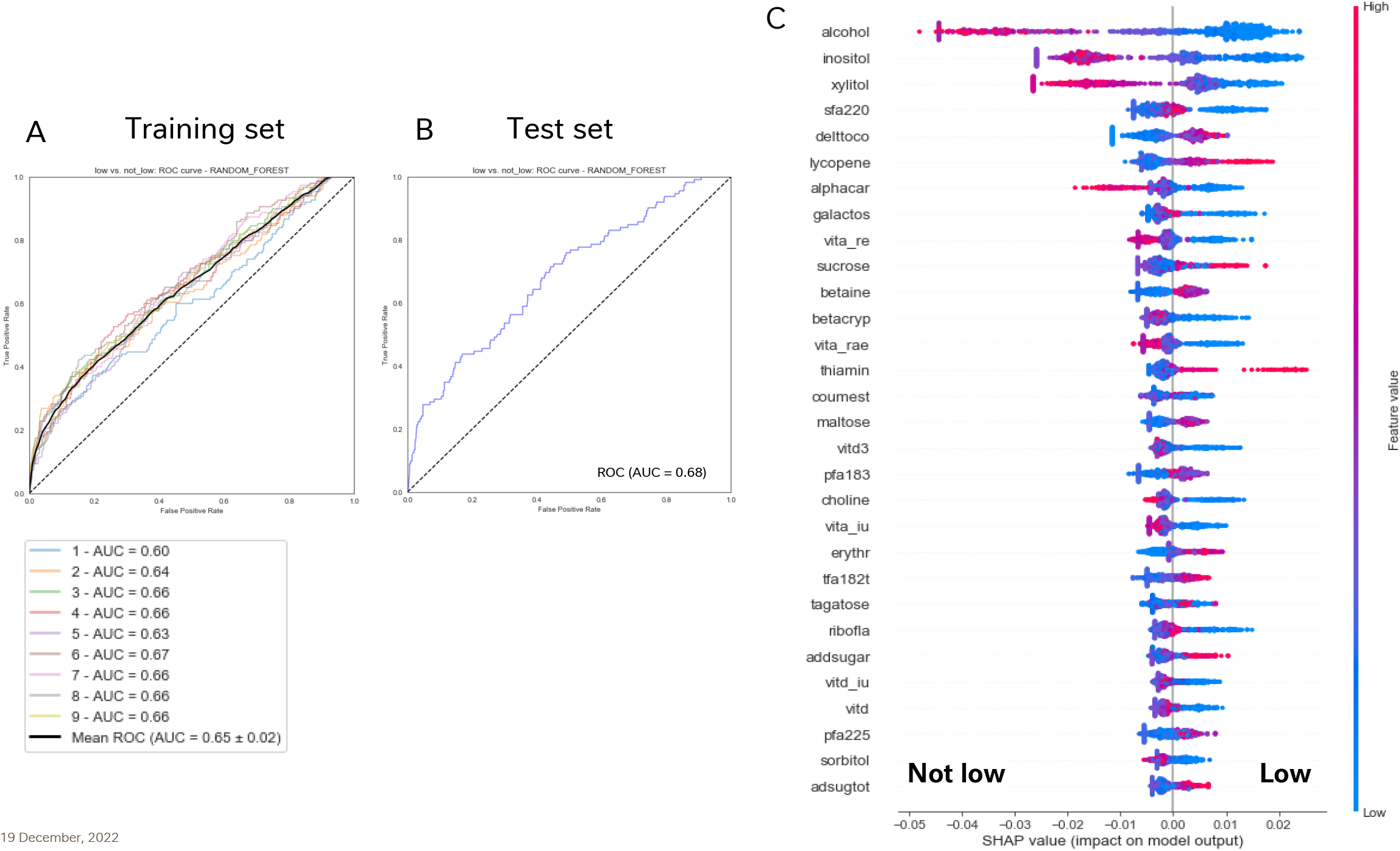
Prediction of relative abundance of *F. prausnitzii* from nutrient intake data. A random forest model was used to predict the individual’s relative abundance of *F. prausnitzii* in low or notlow category (< mean – 1SD vs. the rest). ROC curves and Area Under the Curve (AUC) values for the training (in cross validation mode with 3-folds, 3-repeats) (A) and test (B) datasets are shown. Top 30 most important nutrient features of the model are shown in (C), where contributions of each nutrient feature to the model was analyzed by SHAP [37]. Each dot represents an individual sample. Red and blue color gradations depict higher and lower value of the nutrient’s intake, respectively. If a feature has red values towards the right of the vertical line at 0.00, this indicates higher values of this feature contribute towards “Low” model output. Vice versa, if a feature has red values towards the left of the vertical line at 0.00, this indicates higher values of this feature contribute towards “notLow” model output.

Using agnostic technique SHapley Additive exPlanations (SHAP) [37] to explain predictions of the model, we identified a positive contributions of inositol, xylitol, saturated fatty acid 22:0, a-carotene, galactose and vitamin A to the abundance of *F. prausnitzii* whereas d-tocopherol, lycopene, sucrose and betaine displayed a negative relationship (Figure S4A-S4L).

To complement the above results, we performed univariate analysis (Kruskal-Wallis test) to compare nutrient intakes between the population split according to *F. prausnitzii* relative abundance being Low or notLow, using the same definition of the bins as the best abovementioned model. A total of 11 nutrients were significant after passing false discovery rate (Wilcoxon rank sum test, p.adj<0.05, e.g., alcohol, inositol, aspartame, beta-cryptoxanthin (betacryp), beta-carotene (betacar), total vitamin A activity International Units (vita_iu), total vitamin A activity Retinol Equivalents (vita_re), alpha-carotene (alphacar), pectins, total vitamin A activity Retinol Activity Equivalents (vita-rae) and lutein + zeaxanthin (lutzeax) (Table S3)). When comparing the two *F. prausnitzii* groups (low and notlow) with nutrient intakes normalized to 2000 kcal, alcohol, inositol, aspartame, betacryp and alphcar remained significantly different (Table S4, Wilcoxon rank sum test, p.adj<0.05).

### Growth of F. prausnitzii on inositol-based media is strain dependent

Out of the top nutrients featured in the above analyses, few have previously been shown to support the growth of *F. prausnitzii* in a culture condition, namely: sucrose, maltose and galactose [28, 38]. We therefore selected some of those nutrients to test their potential to enhance *F. prausnitzii* growth *in vitro*, namely: carbocyclic sugar (i.e. inositol) and sugar alcohols (i.e. xylitol, and sorbitol; Figure 1C and Figure S4A, S4B and S4K). Briefly, we measured the growth of two strains of *F. prausnitzii* 27786 and A2-165 representing different phylogenic groups of the bacteria [39]for a period of 48 to 72h on either inositol, xylitol, erythritol or sorbitol as primary carbon source in a YCFA media. We observed that growth under the various tested conditions was strain dependent. Growth of *F. prausnitzii* A2-165 on media prepared with sorbitol was comparable to that observed with glucose as the most efficient carbon substrates followed by inositol and erythritol (Figure 2A) and was further diminished with xylitol to a level close to that with basic YCFA media without any carbon substrate (p=0.0581). In contrast, ATCC 27768 stain grew equally on glucose, erythritol and sorbitol equally, while inositol and xylitol failed to support its growth (Figure 2B).

**Figure 2.**
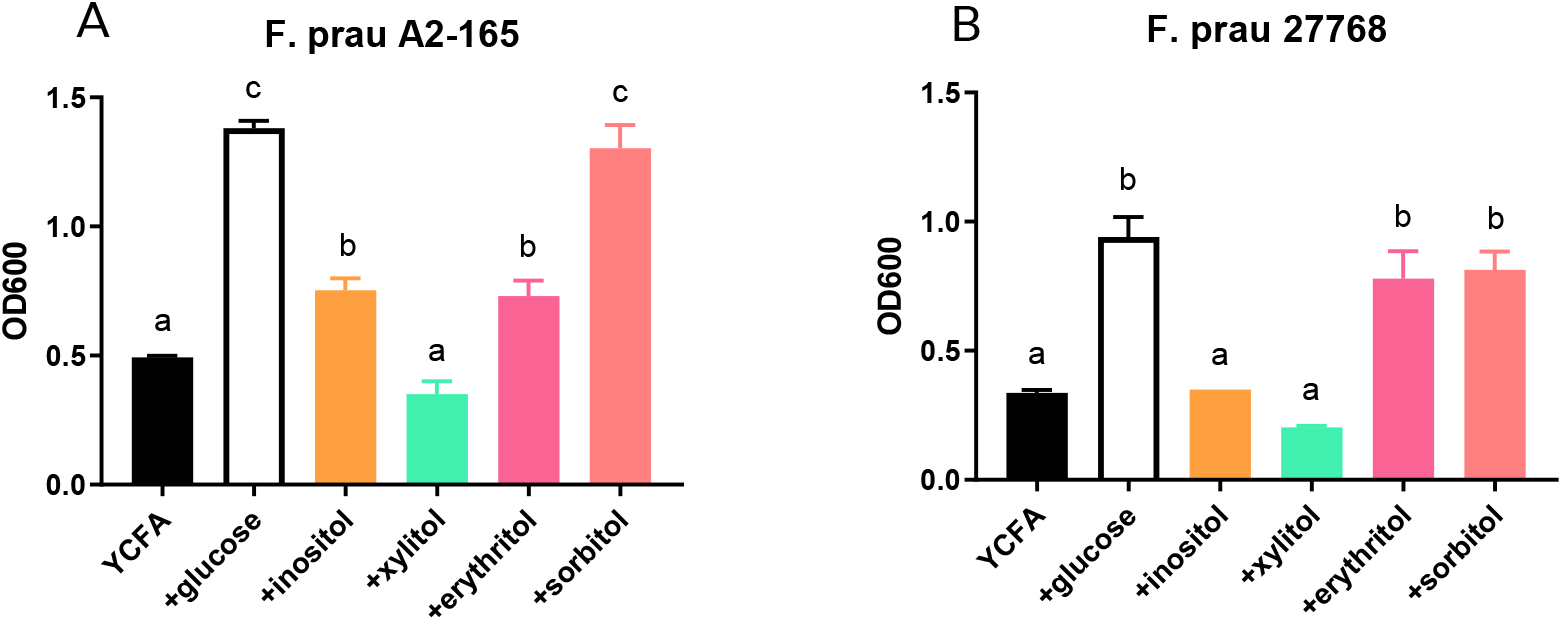
Growth promoting effects of sugar alcohols in two strain of *F. prausnitzii*. Experiments were performed with DSMZ A2-165 (A) or ATCC 27768 (B) in a YCFA based media supplemented with different carbon substrate for 48h in an anaerobic condition. Results are mean +/- SEM, n=3. Statistical analysis was performed with ANOVA followed by Tukey post hoc analysis. Different letter indicates statistical significance between the groups.

We next investigated whether *F. prausnitzii* responds differently with increasing amount of inositol or in combination with other carbon sources. On the inositol-based YCFA media, the growth of A2-165 and 27768 strains only marginally increased compared with YCFA alone (Figure 3A and 3B, p=0.0006 for A2-165 and p=0.0004 for 27768). Doubling the amount of inositol in the media led to 55% more growth with A2-165 strain (Figure 3A, p < 0.0001) but not with 27768 strain (Figure 3B, p=0.7151) when compared with normal amount of inositol. To further illustrate the strain specific substrate utilization, combination of glucose with inositol also increased the growth of strain A2-165 by 21.4% (p<0.0001) compared to glucose alone, while the combination slightly reduced the growth of 27768 by 6.3% (p<0.0001). Finally, addition of inositol to sorbitol promoted growth of A2-165 strain by 64.4% and 23.7% compared to sorbitol alone (p <0.0001), while only a minimal effect of 6.1% was observed on strain 27768 (p <0.0001; Figure 3C and 3D).

**Figure 3.**
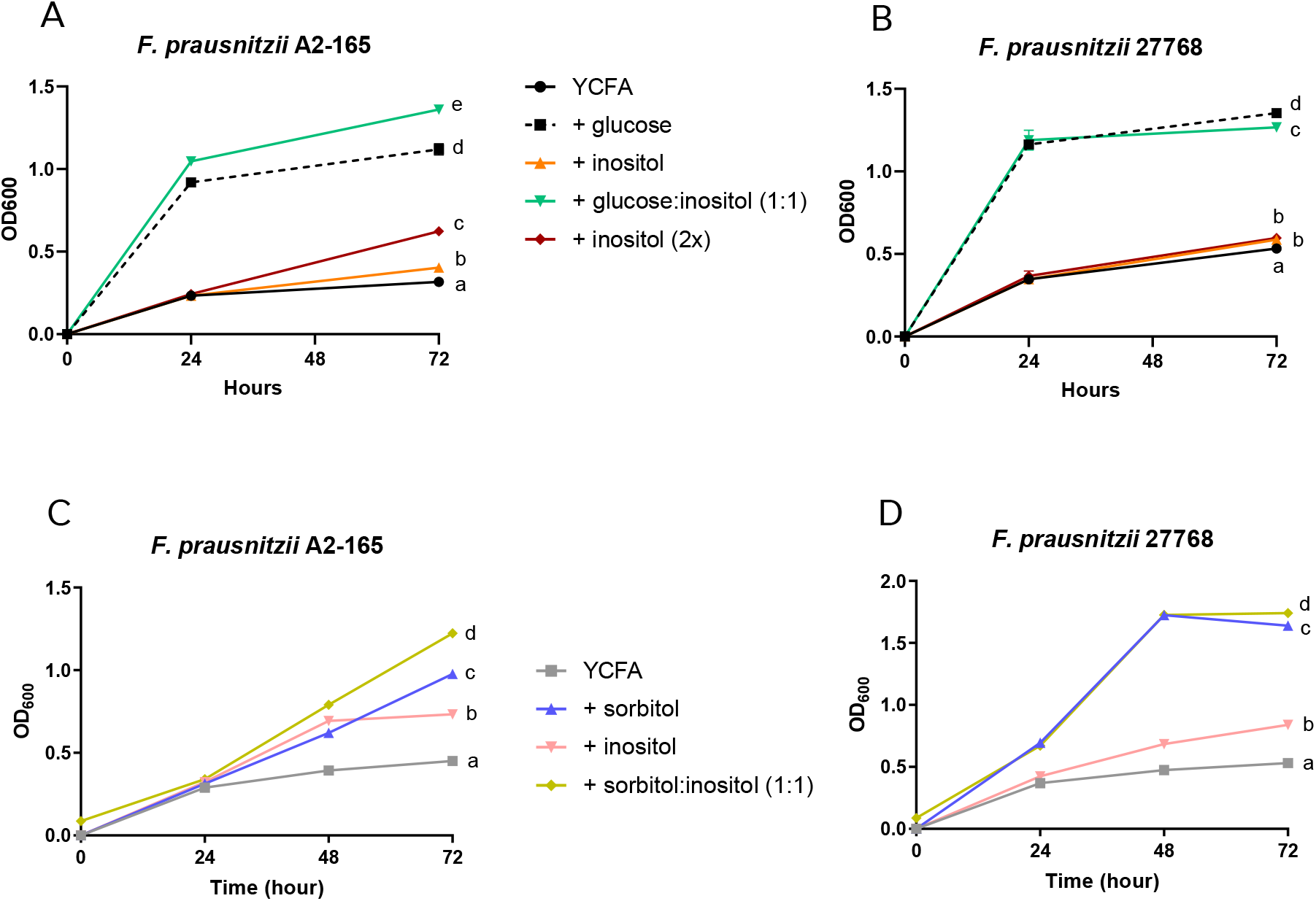
Strain dependent growth response to a combination of nutrients. Growth of *F. prausnitzii* in culture with single nutrient or in combination was followed for 72h and optical density was measure in every 24 h. Glucose (G), inositol (I), glucose and inositol (G+I) and twice amount of inositol (2I) were added to a YCFA media and the results of growth for DSMZ A2-165 and ATCC 27768 are reported in (A) and (B), respectively. Sorbitol (S), inositol (I) or a combination of sorbitol and inositol at equal amount (1:1) were tested in DSMZ A2-165 (C) and ATCC 27768 (D) strains for 72 h. Results are mean +/- SEM, n=3. Statistical analysis of 72h data was performed with ANOVA followed by Tukey post hoc analysis. Different letter indicates statistical significance between the groups.

To further support the predictive potential of the machine learning approach, we also tested whether nutrients predicted to have a negative impact on *F. prausnitzii* would have similar effects experimentally. Results showed that lycopene significantly suppressed the growth of A2-165 by 31.4% (p = 0.039), especially with glucose as the main carbon source (Figure S5A) while betaine failed to alter the growth patten of the A2-165 strain (Figure S5B).

### Responses of F. prausnitzii to nutrients in a mixed community

In a mixed community such as the human gut microbiota, *F. prausnitzii* may compete or work synergistically with other species for nutrients. Hence, response of *F. prausnitzii* to nutrients may highly depend on an ecological context, which could explain discrepancy between the model predictions and the *in vitro* observations described above. Therefore, we tested the effects of nutrient supplementation on *F. prausnitzii* growth by quantitative PCR (qPCR) in an *in vitro* fermentation system with adult human stool samples. Inositol was chosen as the main energy source instead of sorbitol because sorbitol has not been shown to affect the composition of gut microbiota [40] and inositol consistently differentiated *F. prausnitzii* categories in machine learning and univariate analyses. Inulin was used as positive control based on previous reports of a positive effect on the growth of *F. prausnitzii* [38]. In addition, we included vitamin A and E that were identified in the models and B vitamins (B5, B6 and B12) that were not only predicted by the model but also essential for *F. prausnitzii* [41].

Effects of nutrients on *F. prausnitzii* growth were tested for a period of 48h using fecal samples from 4 individual donors as replicates in a casitone-based oligotrophic media. With most interventions, we observed a non-significant increase of *F. prausnitzii* compared to control, especially after 24h (Figure 4A and 4B). A high degree of heterogeneity in the response was observed across fecal donors (Figure S6A-S6D). For instance, treatment with inulin resulted in a 24.5 and 10.6 folds increase in *F. prausnitzii* at 24h compared to control in donor 2 (D2) and 3 (D3) respectively, while no effects were observed with donor 1 (D1) and 4 (D4). Inositol alone or inositol with vitamin supplementations also triggered an increase in *F. prausnitzii* by at least 50% compared to control in D2 and D3 communities, and yet no effects were observed with D1 and D4 (Figure S6A-S6D).

**Figure 4.**
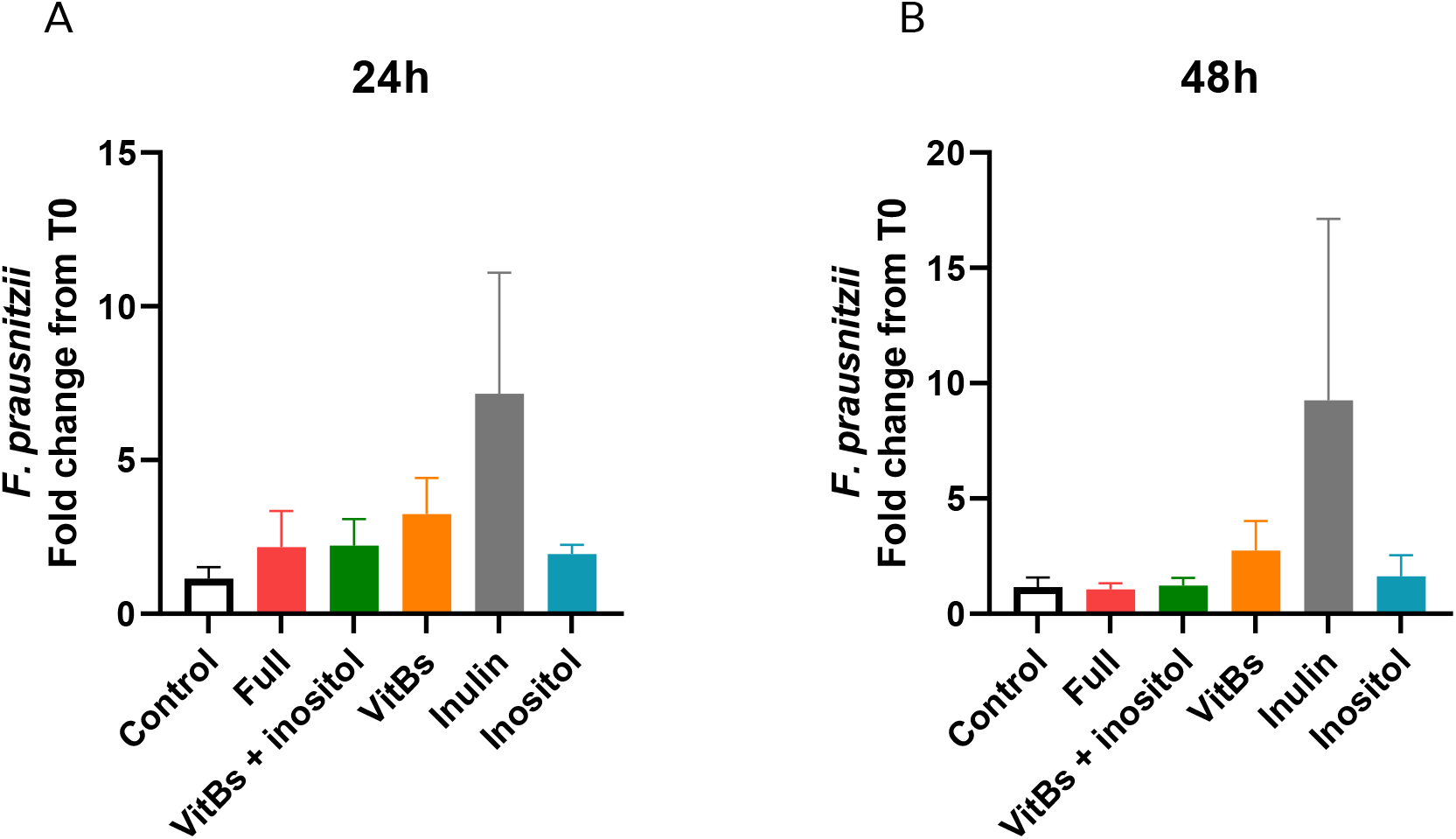
Evaluation of nutrients and nutrient combinations on the growth of *F. prausnitzii* in a mix community. Details of *in vitro* fermentation experiments are described in Methods and Materials. The fold changes of *F. prausnitzii* at 24h (A) and 48h (B) from the baseline were calculated from results of F. prausnitzii specific qPCR quantification. Each treatment was tested in 4 fecal samples and data are shown in mean +/- sem. Statistical analysis was performed with ANOVA followed by Tukey post hoc analysis, but no statistically significant difference was found.

To further understand the heterogeneity of these results, we next conducted metabolomics profiling of the fermented media at all time points. PCA with 34 identified and integrated NMR signals of metabolites suggested that time had more effects on the metabolomic variance during the fermentation than donor or treatment. While the 6h time points clustered closely with the baseline samples, a drastic change in overall metabolic profile was observed at 24 and 48h (Figure S7A and S7B). PCA loadings showed that from 6 h to 24 h of fermentation, short-chain fatty acids, trimethylamine, alcohols, monamine aromatic amino acid-derivatives, diamines and related metabolites increased, while glycerol and some amino acids (threonine, tryptophan, tyrosine and arginine) were decreased (Figure S7C). Lactate, formate and succinate increased over 24 h before being consumed at 48 h.

Focusing more precisely on the metabolization of the tested substrates, we observed in all four donors, that inositol was fully consumed over time (Figure S8A to S8C) independently of vitamin supplementation, and the rate of consumption was not related to level of *F. prausnitzii*. Finally, as *F. prausnitzii* is one of key butyrate producing bacteria [42], we examined the correlation between *F. prausnitzii* and butyrate in the batch fermenters. Taking all samples into account, we observed a positive correlation between levels of *F. prausnitzii* and butyrate concentration (Pearson correlation, r=0.6252, p<0.0001; Figure 5). Even after removing the leverage point, result is still significant (p<0.0001, r=0.4661). This correlation remained significant when considering D3 alone while only a trend was observed for D1 and D4 (Figure S9).

**Figure 5.**
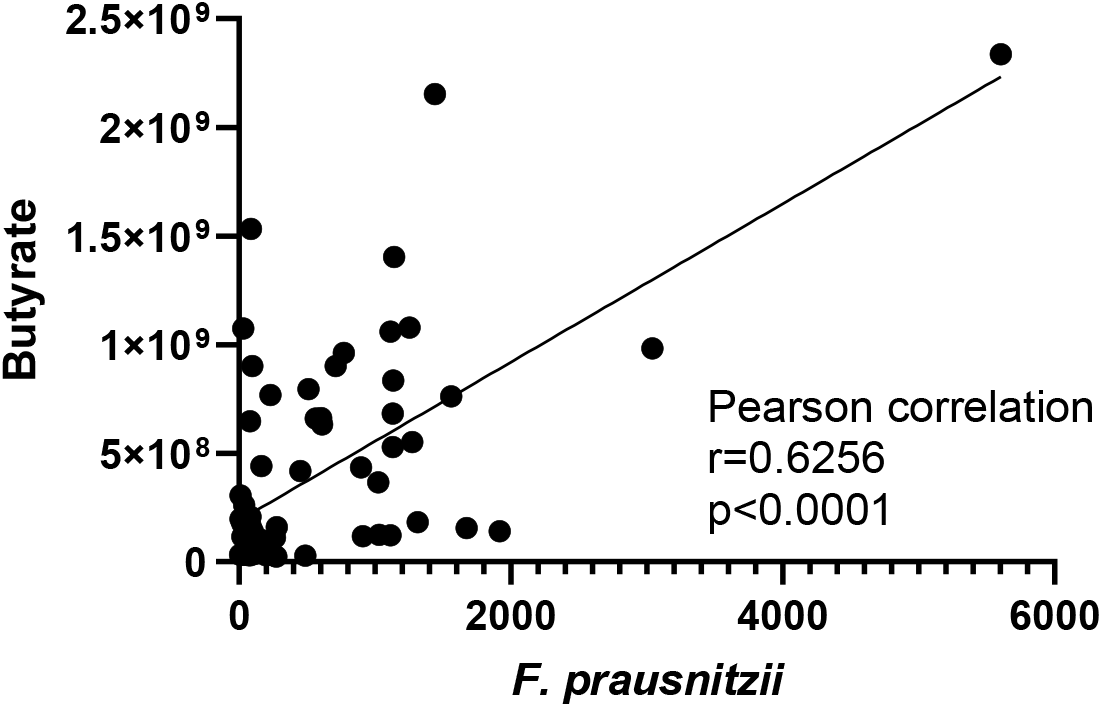
Relationship between butyrate and the amount of *F. prausnitzii* in batch fermentation experiments. Signals of butyrate was extracted from ^1^HNMR metabolomics and the amount of *F. prausnitzii* was determined with qPCR technique. Data were pooled from 6, 24 and 48 h time points with all treatment groups in all four donors. Pearson correlation analysis indicates a positive relationship between the number of *F. prausnitzii* and butyrate (p<0.0001, r=0.6256).

## Discussion

*F prausnitzii* is amongst the most abundant anaerobic bacteria in human gut, and scientific evidence supports its beneficial role in health. In the present study, we applied a machine learning algorithm to microbiome composition and food frequency questionnaires data collected on 3816 AGP participants and identified nutrients that may influence the abundance of *F. praustnizii*. Subsequent *in vitro* experiments with two strains of *F. prausntizii* demonstrated that inositol, sorbitol and lycopene could enhance the growth of at least one of the selected bacterial strains as predicted by the model. On the contrary, xylitol, erythritol and betaine failed to increase *F. prausnitzii* growth under *in vitro* conditions suggesting that other factors than these nutrients alone may be at play. More importantly, we observed strain-dependent responses of *F. prausnitzii* to most nutrients or nutrient combinations. Besides, when the effects of nutrients on *F. prausnitzii* were tested in the context of complex communities using *in vitro* fermentation, we observed a high degree of variations among the four fecal donors, rendering no significant changes in the number of *F. prausnitzii*. Interestingly, we observed a significant positive correlation between *F. prausnitzii* and butyrate concentration during fermentation, supporting the use of *in vitro* fermentation models to study microbial metabolism.

A citizen science project such as AGP offers a large dataset for examining the relationships between gut microbiota and a wide variety of factors such as dietary patterns, lifestyle, diseases, etc. [24, 44-47]. We included previously un-processed 16S composition data and created a cohort of 3816 AGP subjects for this study. Compared to typical American adult population (NHANES), the study cohort seemed to consume less calories and have healthier eating habits for children, adults, and to a lesser extent for older adults. Contrary to calorie intake, AGP cohort reported a higher intake of fiber and vitamin B12 than NHANES, further supporting a healthy eating choice of AGP participants. It is however worthwhile to mention that different methods of collecting dietary intake in the two studies hindered us from performing direct comparison between the two cohorts, similar to the conclusion of a recent study looking at dietary pattern of 1800 AGP participants [45].

Through our modelling approach employed here, many nutrients identified in this study were newly associated with *F. prausnitzii*, such as sorbitol and inositol, while others such as alcohol and galactose have been previously reported to positively correlate with high *F. prausnitzii* abundance[32, 48].

Inositol or myo-inositol is commonly found in vegetables and meat [49], and sorbitol and many sugar alcohols are found in fruits and vegetables. Evidence from a prospective study and a dietary intervention study showed a positive relationship between the consumption of fruits and vegetables and the abundance of *F. prausnitzii* [50, 51], a result in agreement with the findings of our *in vitro* experiments. On the other hand, we did not observe any growth promoting effect of xylitol on either strain of *F. prausnitzii*. These results highlight the importance of experimental validations on the outcomes of *in silico* modeling.

*F. prausnitzii* has a high degree of genetic diversity, and the two strains used in the study (A2-165 and 27768) belong to different phylogroups [38]. The two isolates also showed different growth rate under various dietary and host derived carbohydrate sources [38]. We also observed strain specific growth response when inositol was given as a carbon source. Recently, a branch of *F. prausnitzii*, including A2-165 strain has been reannotated into a new species *Faecalibacterium duncaniae* [52], further highlighting the diverse metabolic potential of *F. prausnitzii*. Interestingly, it was reported that neither *F. duncaniae* nor *F. prausnitzii* grew on inositol, which contrasts with our findings. The discrepancy is likely due to the difference in culture condition: in our study, the growth promoting effect of inositol was observed after 48 h whereas Sakamoto et al. [52] reported results after 18-24 h incubation time.

Use of *in vitro* fermentation in a test tube has been widely applied to examine microbial degradation and transformation of prebiotic fibers [53] due to many advantages such as short turnaround time, enhanced throughput, simple equipment set up compared to a continuous system, animal models or clinical studies [30]. However, it is also the least physiological of all the models as pH is often not fully controlled and waste products are not removed during the fermentation. Despite the shortcomings of the system, we showed that inositol was efficiently utilized by all four fecal communities, and *F. prausnitzii* increased at least 1.6-fold over control in three out four communities. One reason contributing to the interindividual variation could be the differences in microbial composition among all the donor samples. *F. prausnitzii* is highly connected with other bacterial members in the energic trophic chain. This is best demonstrated in cross-feeding experiments where *F. prausnitzii* population benefited from the presence of *Bifidobacteria* and other bacteria for acetate and vitamin Bs, respectively [54, 55]. Since *Bifidobacteria* are primary utilizers of inulin in adult gut ecosystem [56, 57], it is not a surprise to see donor-specific responses to test nutrients and inulin. *F. prausnitzii* also completes with other bacteria for carbon sources. Shown by Lopez-Siles et al., *F. prausnitzii* out-competed *Eubacterium eligens* and *Bacteroides thetaiotaomicron* in co-culture experiments with apple pectin. However, it is possible *F. prausnitzii* does not have competitive advantage for other nutrients. To concretely evaluate the effect of *F. praustnitzii* targeting nutrients, intervention trials in humans coupled with metagenomic and metabolomic analysis are needed to reveal nutrient-*F. prausnitzii* relationship in a complex gut ecosystem.

In conclusion, we discovered novel *F. prausnitzii* modulating nutrients using machine learning approach applied to data from American Gut Project, and many of our predictions were confirmed in *in vitro* experiments, supporting the value of *in silico* approach without having *a priori* hypothesis. One of potential value of our model is that it deals with the lower spectrum of the population-wise distribution of *F. prausnitzii* in AGP, which helps us identify subjects who may benefit from nutritional interventions to boost *F. prausnitzii*. While validating the nutrients singly or in combinations, we experienced highly individualized responses among 4 fecal donors. These findings could serve a foundation for future clinical trial design focusing on personalized nutrition for targeted modulation of beneficial bacteria such as *F. prausnitzii* in humans, to support the maintenance of healthy microbiota in adults.

## Acknowledgements

Many thanks to Lakshmi Rajasekhar, who generated the models and SHAP interpretation plots when employed by Intellify, Australia. We also deeply thank Ms. Shivangi Verma, Mr. Santosh Elavalli, Dr. Palani Kannan Kandavel and Dr. Pramila Tata, formerly of Syngene International Limited, India for processing of 16s rRNA gene sequencing data.

## Patents

Two patents were filed related to the works discussed here:

1. Systems and methods for estimating, from food frequency questionnaire-based nutrients intake data, the relative amounts of Faecalibacterium prausnitzii (Fprau) in the gut microbiome ecosystem and associated recommendations to improve Faecalibacterium prausnitzii https://worldwide.espacenet.com/patent/search/family/075825630/publication/WO2022233924A1?q=pn%3DWO2022233924A1
2. Compositions and methods using at least one of inositol or sorbitol to enhance growth of Faecalibacterium prausnitzii https://worldwide.espacenet.com/patent/search/family/081941092/publication/WO2022233922A1?q=pn%3DWO2022233922A1

## Figure legends

**Figure S1.**
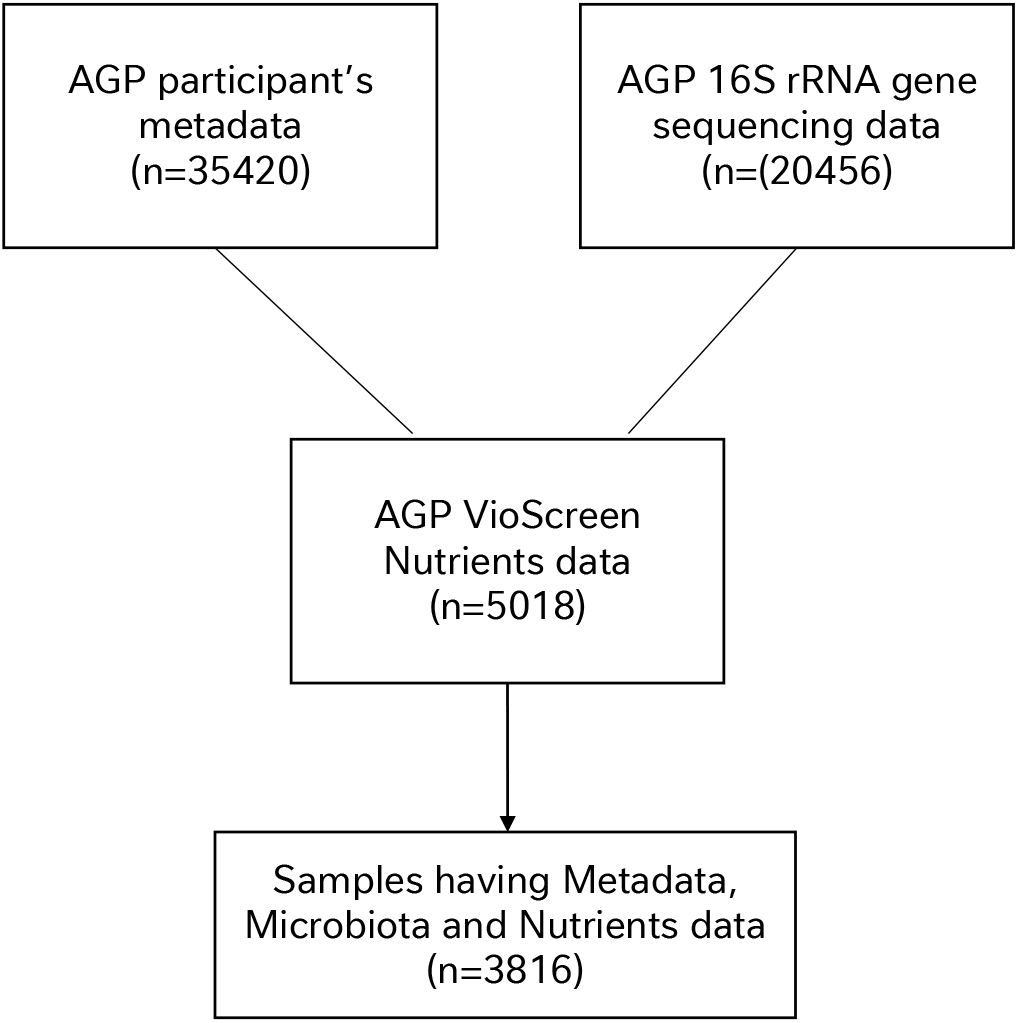
Selection of subjects for the analyses. American Gut Project (AGP) Metadata (n=35,420), 16S rRNA gene sequencing Microbiota data (n=20,456) and FFQ-based Vioscreen-derived Nutrients data (n=5,018) were downloaded from the respective repositories described in Methods and Materials. Only the subjects having all three data types available were included in the modeling analysis (n=3,816).

**Figure S2.**
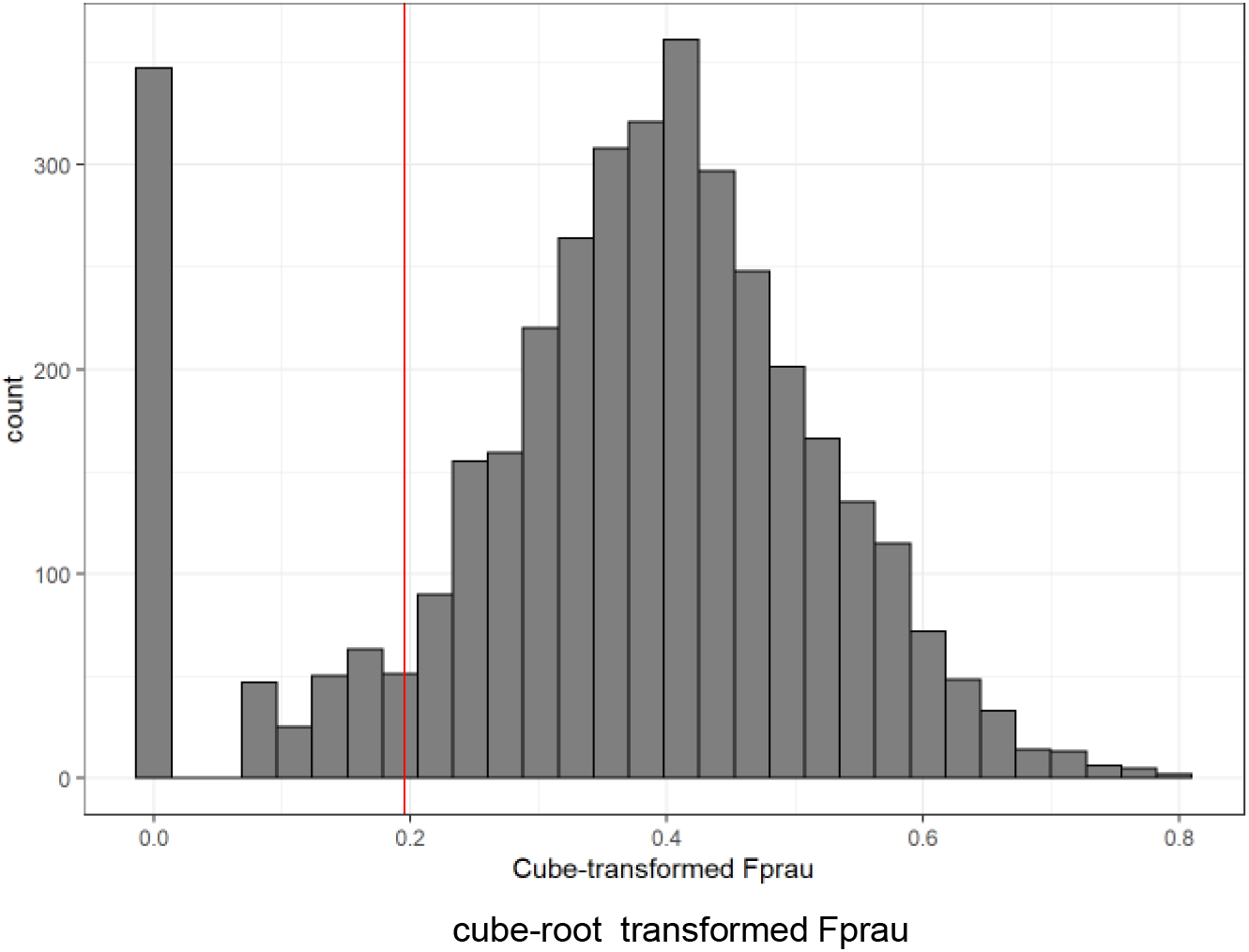
Distribution of *F. prausnitzii* in the AGP sub-cohort. A subset of AGP subjects (n=3816) was selected with available metadata, FFQ data and 16S data. The relative abundance of *F. prausnitzii* was cube-transformed and the frequency distribution is shown. A normalized distribution was then used to create different categories (i.e. low vs notlow, high vs nothigh and low vs high).

**Figure S3.**
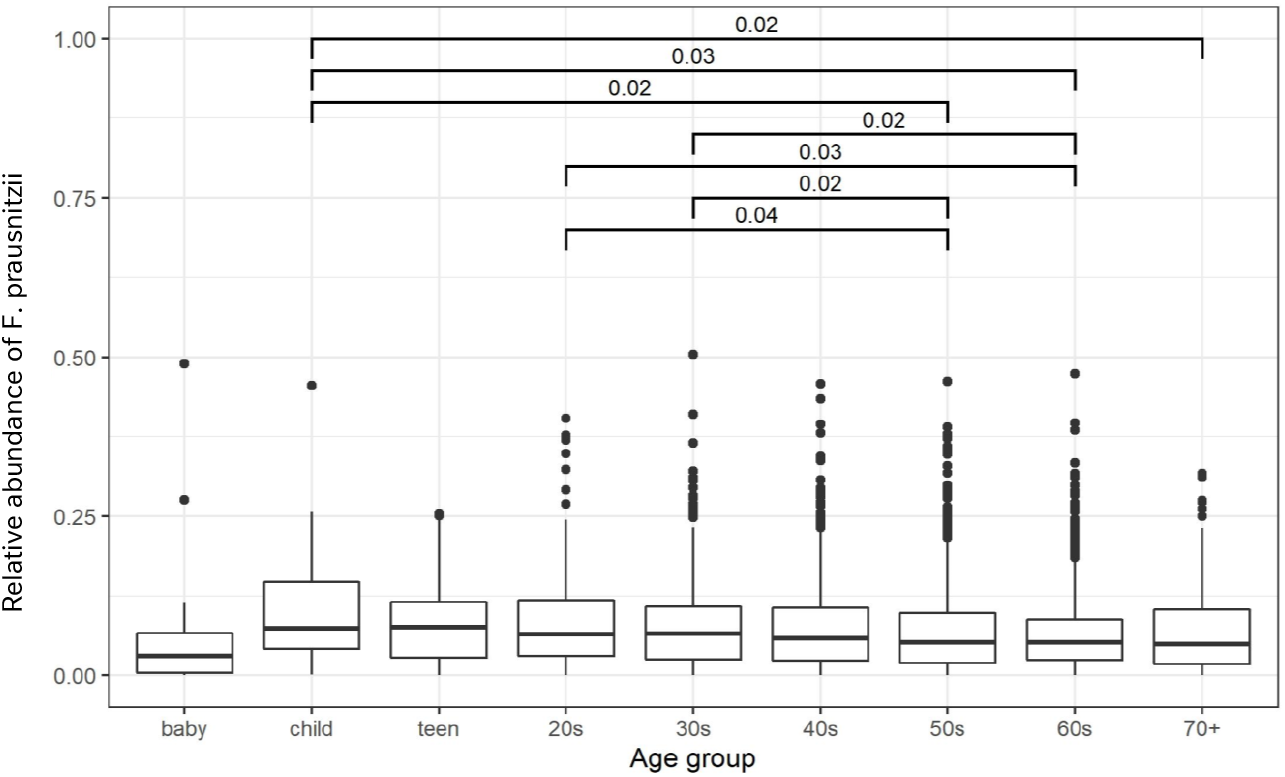
Age-associated relationship with *F. prausnitzii* in AGP study cohort. Relationship between the relative abundance of *F. prausnitzii* and self-reported age of 3816 AGP subjects. Differences in *F. prausnitzii* relative abundance between age categories are shown. Statistical comparisons were performed with Kruskal-Wallis rank sum test followed by post hoc Dunn test. p values of statistically significant differences are shown on the top of cross bars.

**Figure S4.**
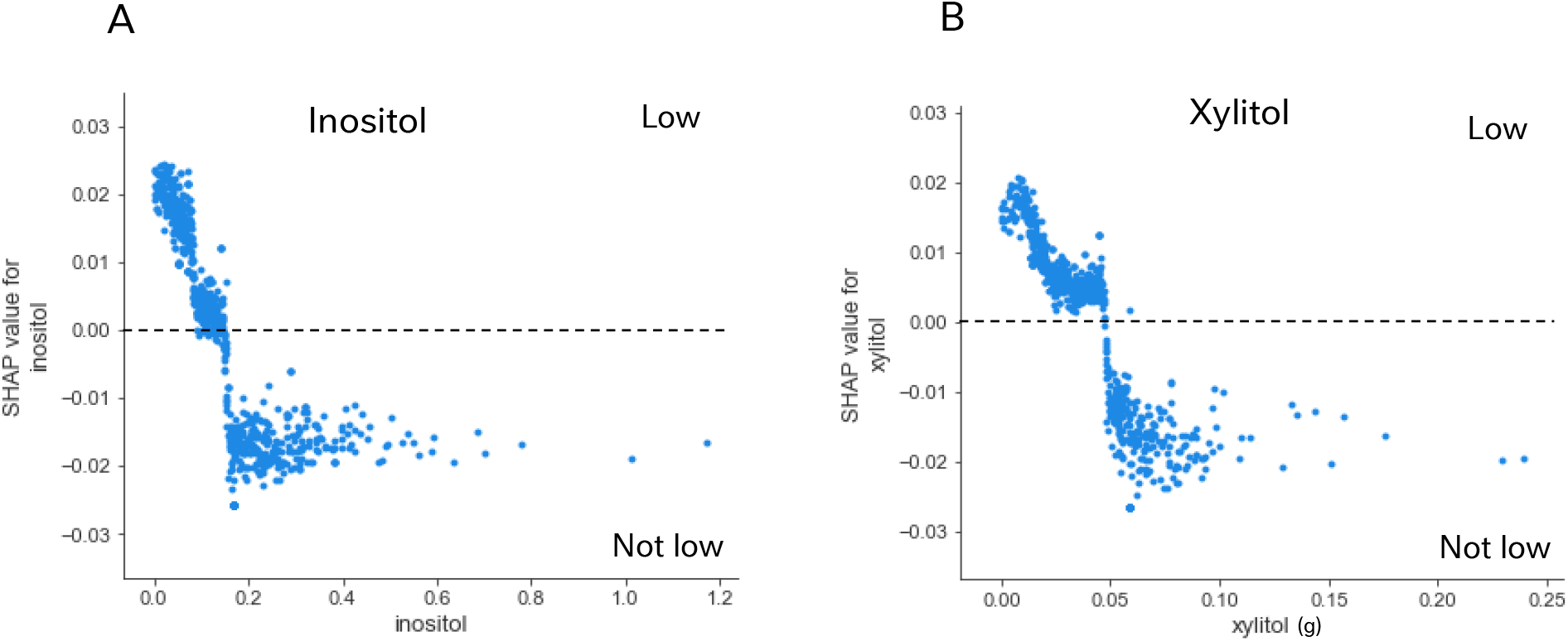
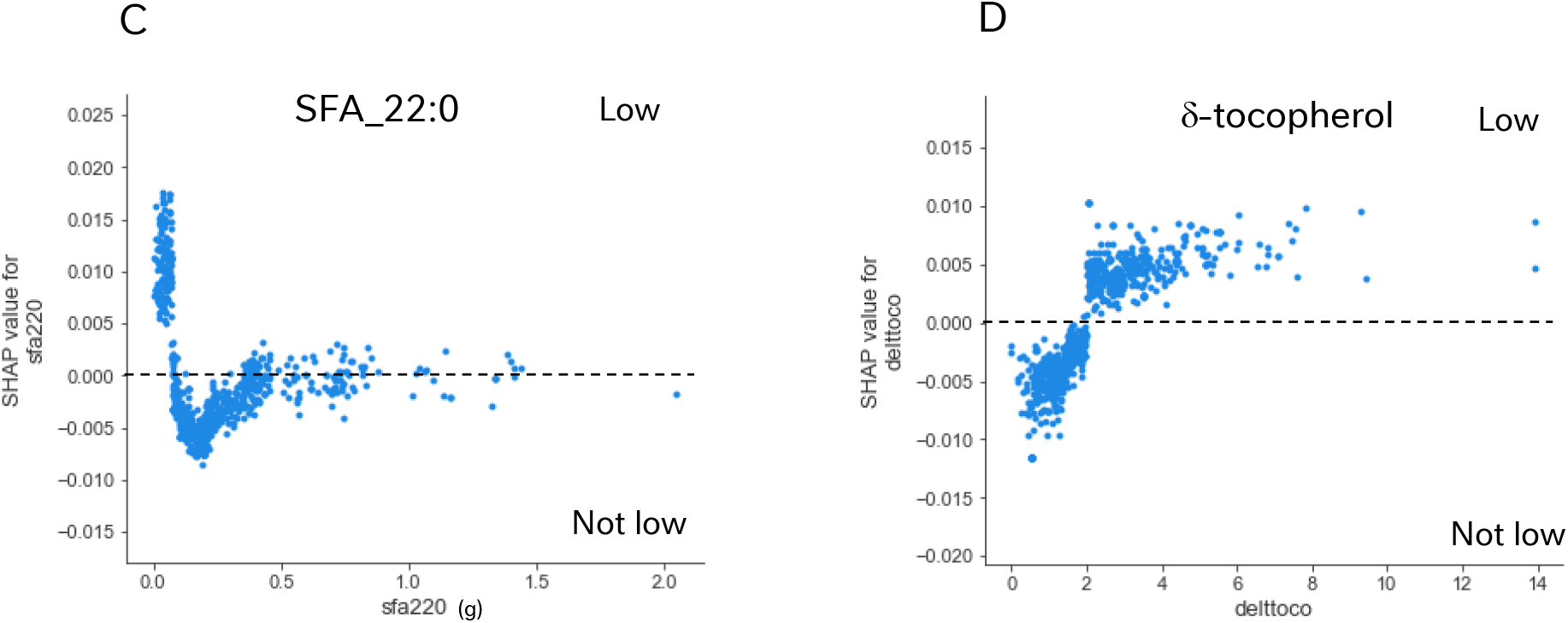
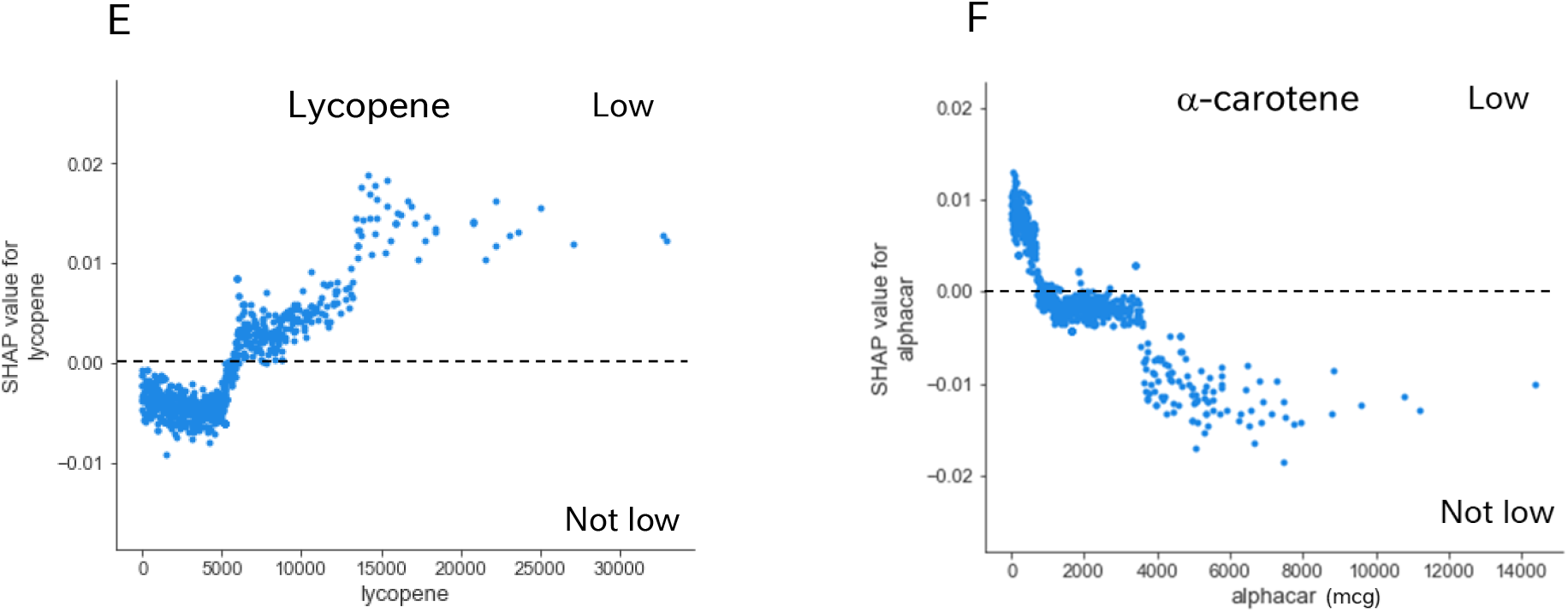
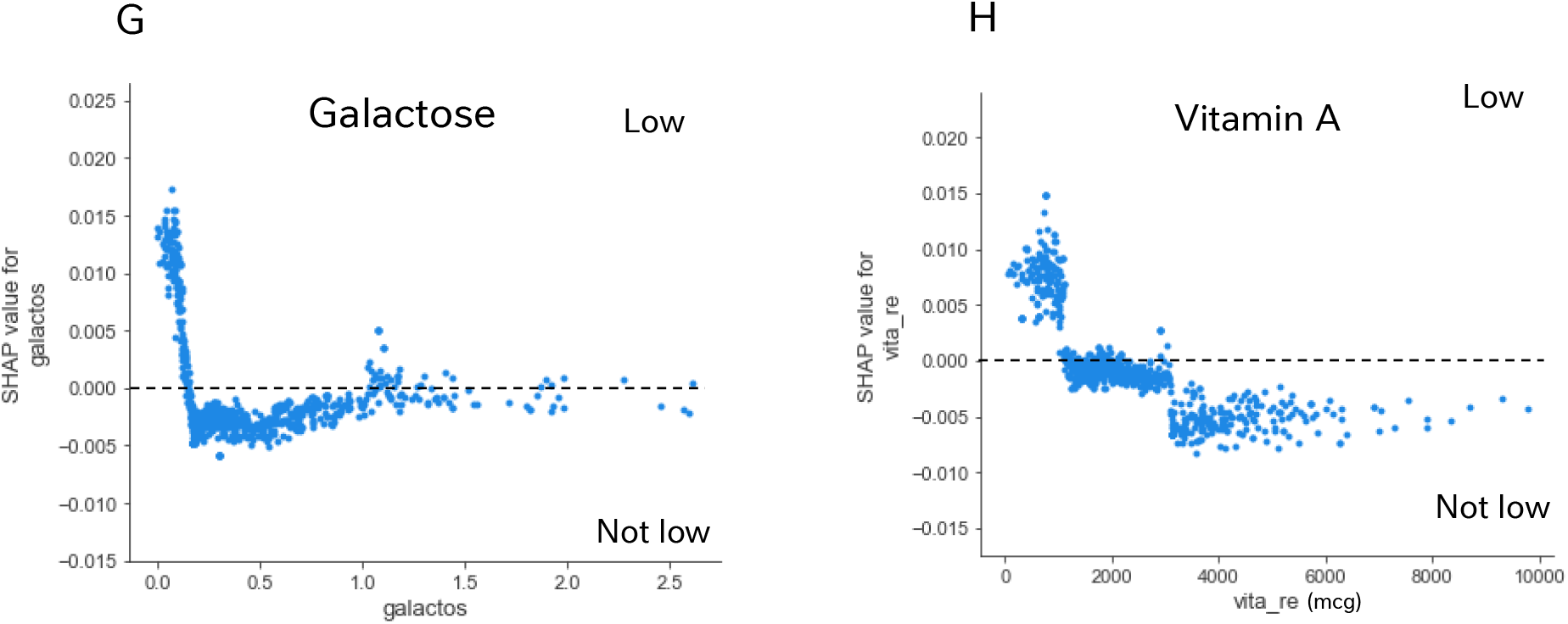
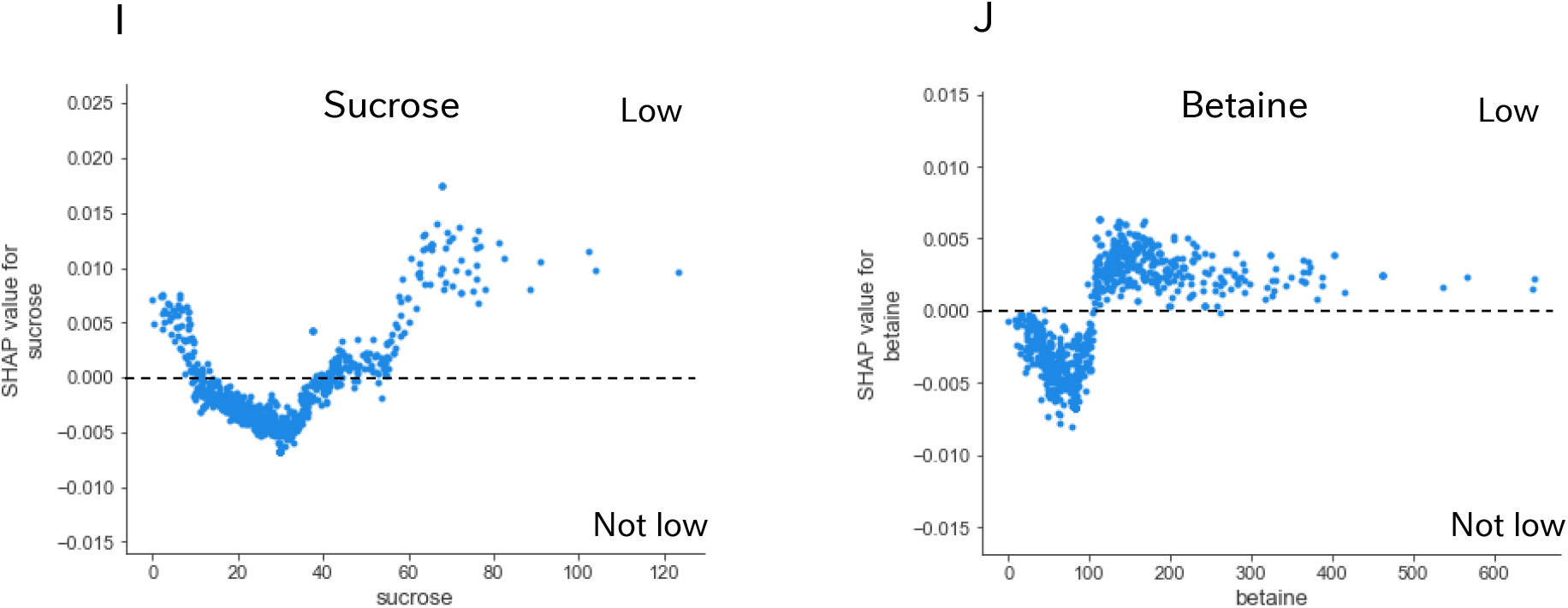
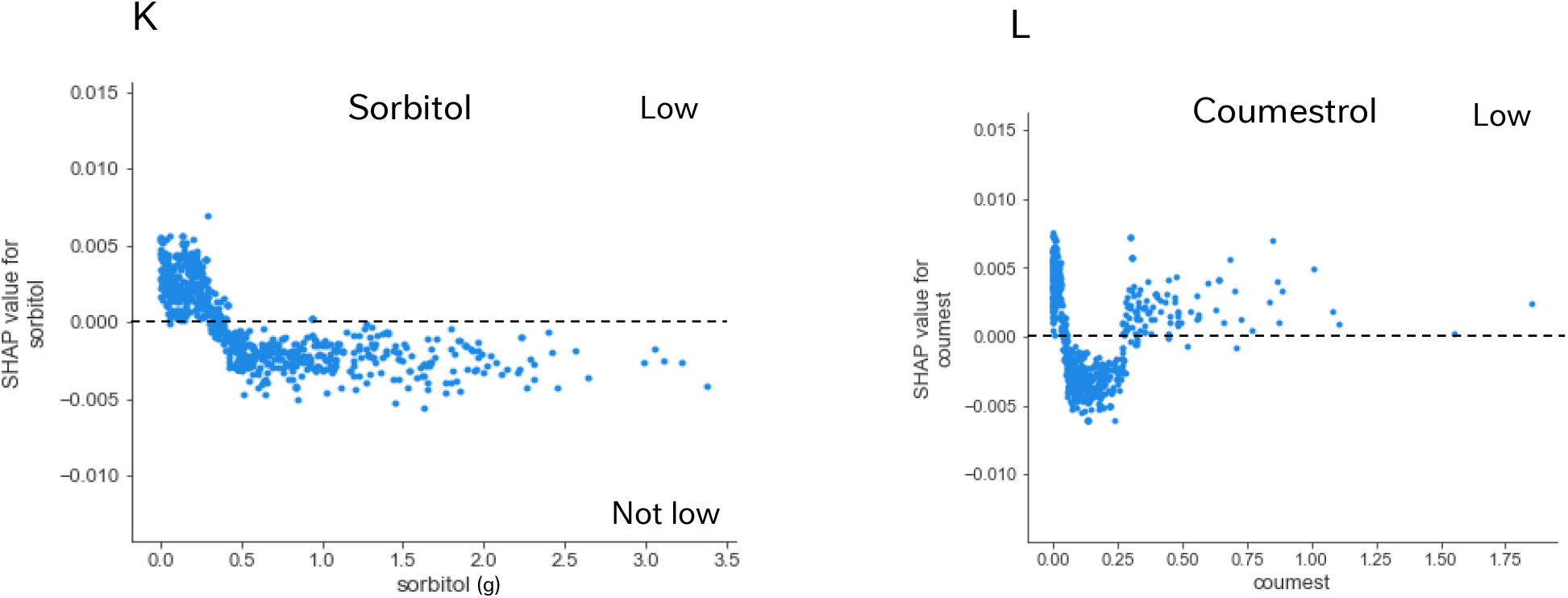
SHAP Dependence plots of top nutrients in the model. SHAP dependence plots of top 12 nutrients except for alcohol are presented in (A, inositol); (B, xylitol); (C, SFA_22:0); (D, d-tocopherol); (E, lycopene); (F, a-carotene); (G, galactose); (H, vita_re); (I, sucrose); (J, betaine); (K, sorbitol); (L, coumest). For each nutrient feature, the SHAP Dependence plot shows the intake value on the x-axis and the corresponding Shapley value on the y-axis. The reference class here was “Low”. Thus, the positive coefficients of SHAP value on the y-axes, with the corresponding x-values, indicate how the model was affected in predicting the “Low” class using this feature. For example, as shown in Supporting figure 1A, specific intake values of inositol had a relation with impact on model output. Low inositol intake has the *F. prausnitzii* status in the “Low” class, while higher intake amounts of inositol have the Fprau status in “notLow” class. When interpreting, please note that the final prediction of the model is result of a complex multivariate analysis. Thus, the final impact on the *F. prausnitzii* status of an individual, e.g. as Low or notLow, was a combination of different features to one single output from the model.

**Figure S5.**
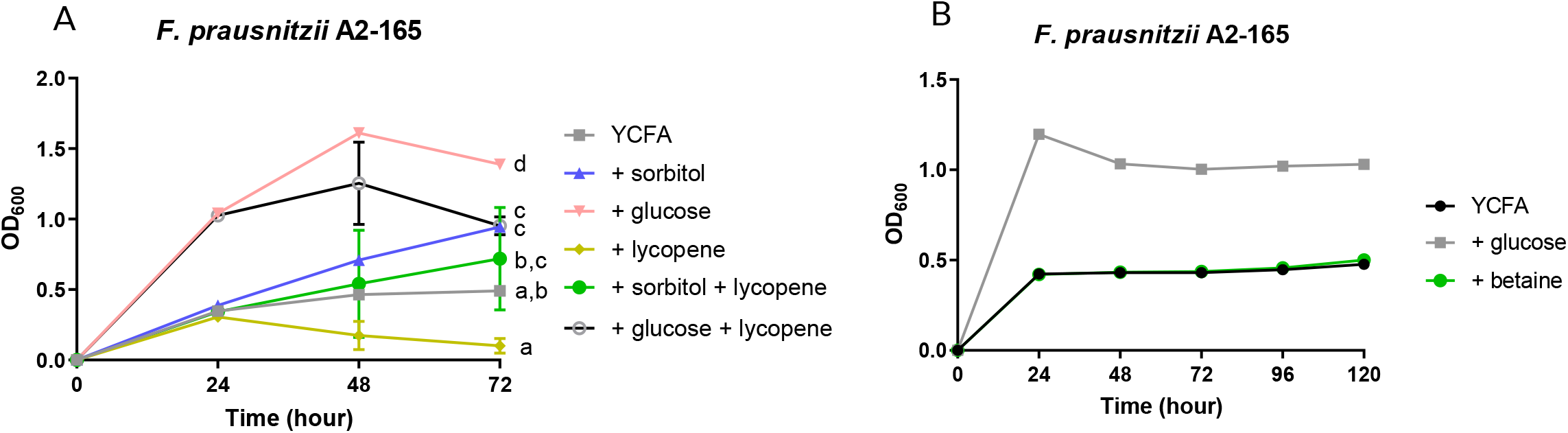
The growth of *F. prausnitzii* in single or combination of nutrients. Anaerobic culture conditions of *F. prausnitzii* A2-165 are described in Methods and Materials. The growth of the bacteria over time was measure with optical density (OD_600_). Response of *F. prausnitzii* to glucose, sorbitol, lycopene, sorbitol+lycopene and glucose+lycopene are shown (A). Growth of the bacteria under glucose or betaine is shown in (B).

**Figure S6.**
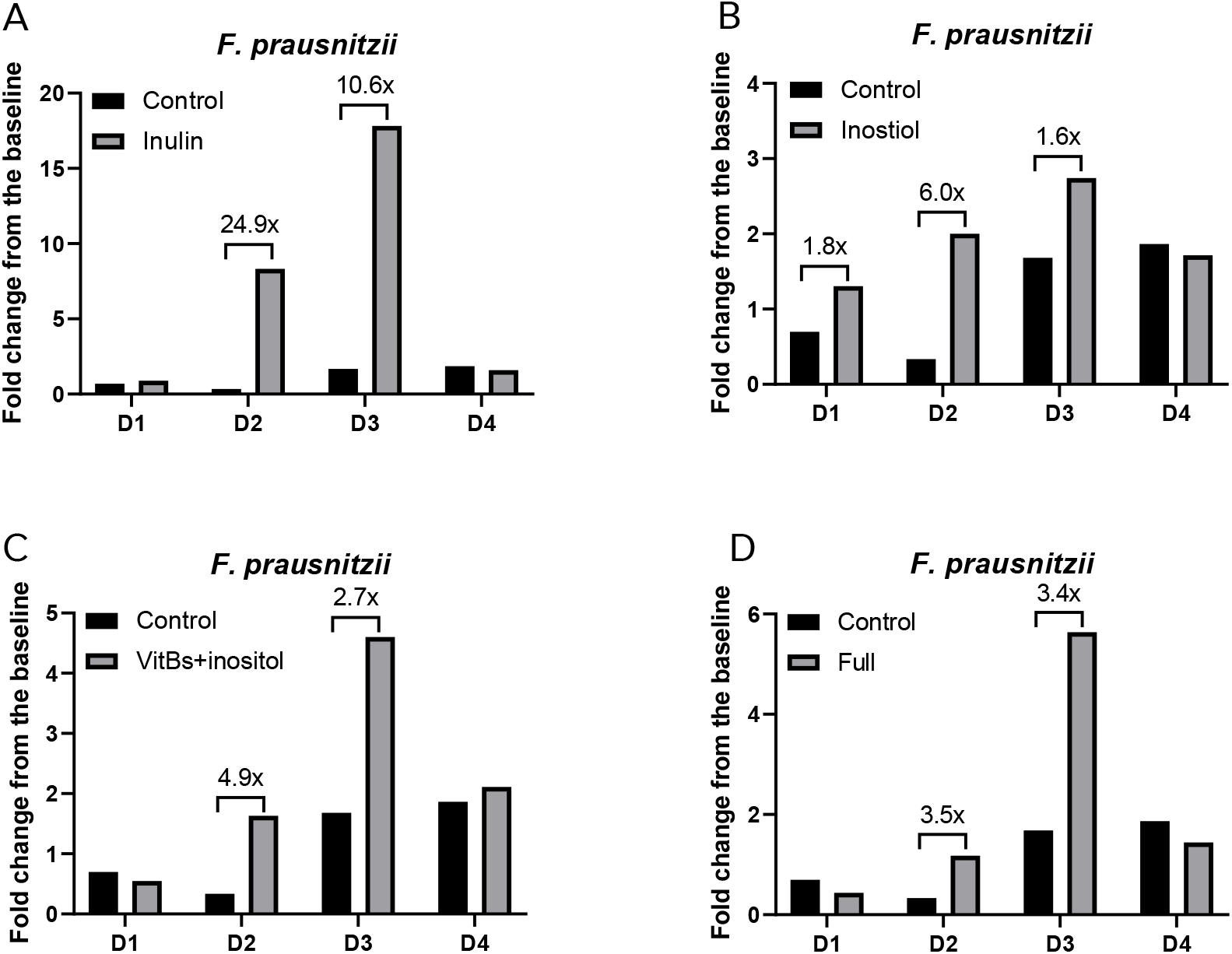
Effect of nutrients on the growth of *F. prausnitzii* in a complex community is donor dependent. The effect of single or combination of nutrients on number of *F. prausnitzii* in a mix community was examined in an in vitro batch fermentation system as described in Methods and Materials. The number of *F. prausnitzii* was quantified by qPCR using *F. pruastnizii* specific primers. The results of inulin (A), inositol (B), vitBs+inositol (C) and full (D) are shown. vitBs consists of vitamin B5, B6 and B12, and full denotes Vitamin A, vitamin B5, vitamin B6, vitamin B12, vitamin D. Fold change of *F. prausnitzii* at 24h from the baseline are shown for each of the four microbiome backgrounds and the differences between treatment and control are indicated when the number is larger than 1.5.

**Figure S7.**
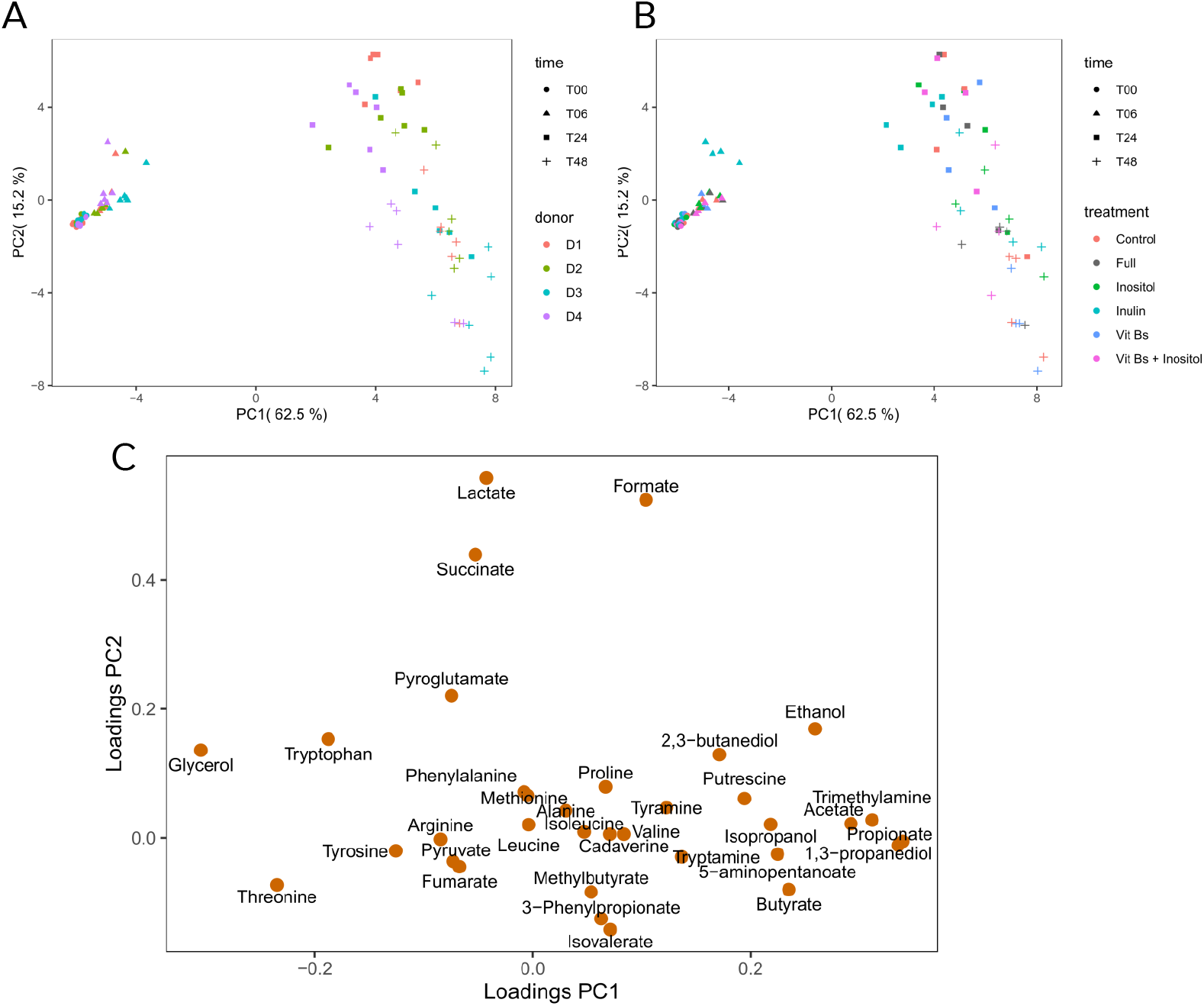
PCA analysis of metabolomic profiles of *in vitro* fermentation media. Effects of nutrient or nutrient combinations were examined in four microbiota backgrounds. Samples were collected at time 0, 6, 24 and 48h from batch fermentation experiments. Each dot in the PCA score plot represents an individual, and the shape and color of symbol denote the time point, fecal donor or nutrition treatment (A and B), respectively. The loading plot highlighted the metabolites influencing distribution in PCA score plots (C).

**Figure S8.**
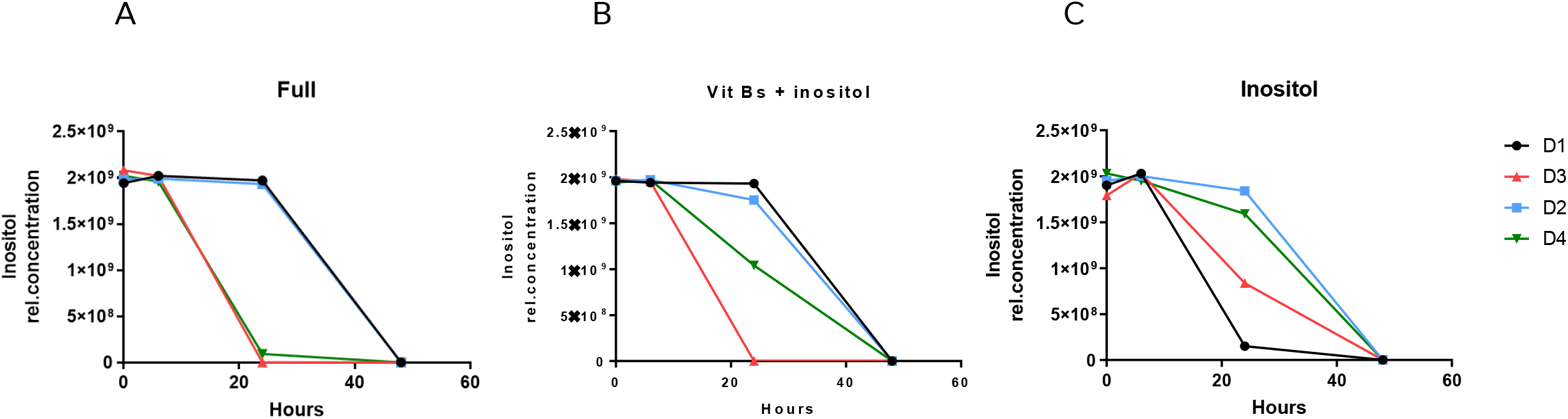
Decrease of inositol signal over the duration of *in vitro* fermentation. Signal of inositol was extracted from metabolomic data and strength of the signal at each time point is shown. Results of full mix (A), vitB+inositol (B) and inositol alone are shown. Color of line represents different microbiota background (D1-D4).

**Figure S9.**
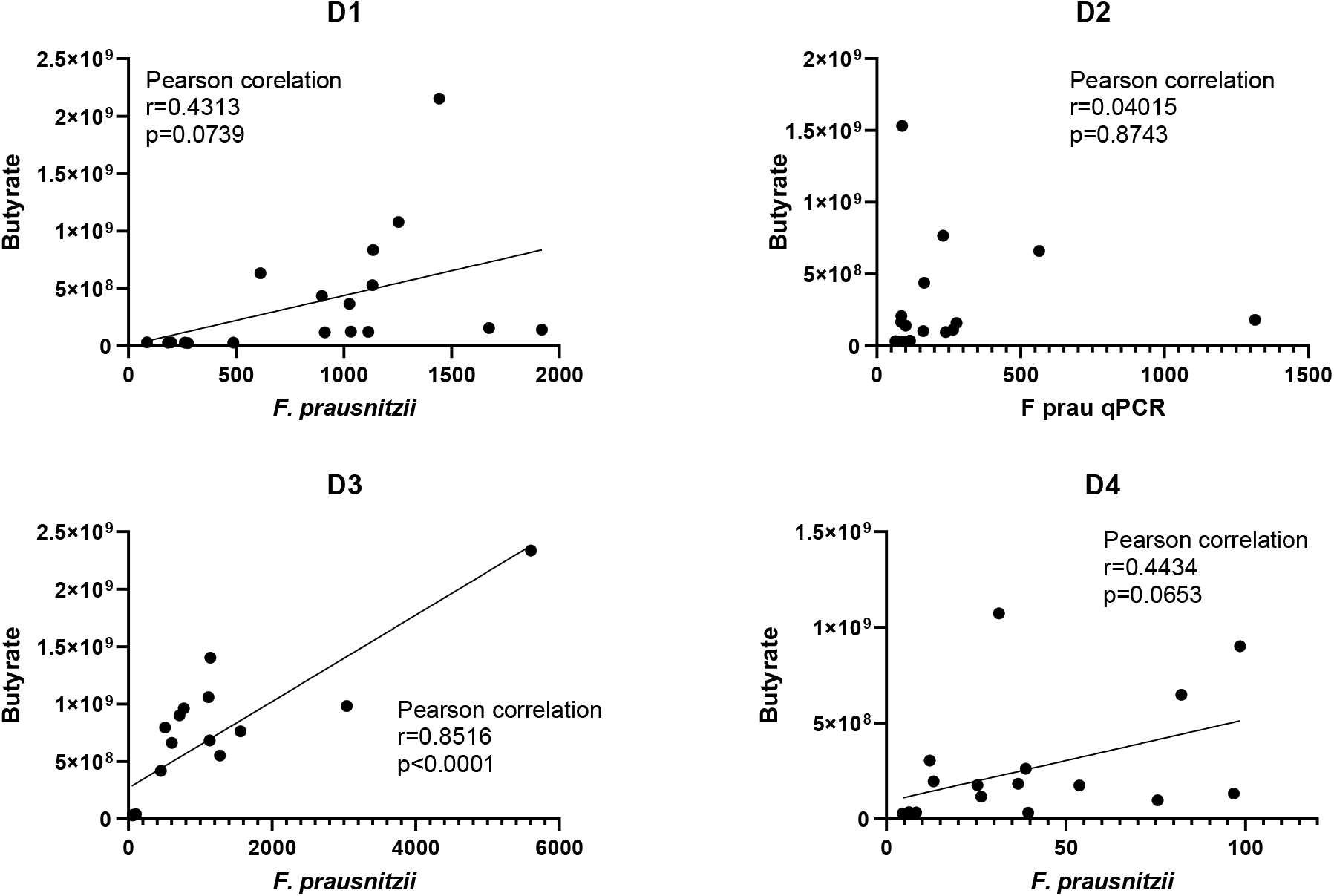
Relationship between the number of *F. prausnitzii* and butyrate signals in the fermentation media. *F. prausnitzii* number was measured with *F. prausnitzii* specific qPCR, and butyrate signals were extracted from ^1^HNMR spectra. Results include all time points (0, 6, 24 and 48h) and are presented according to the microbiota background, donor 1 (A), donor 2 (B), donor 3 (C) and donor 4 (D). Pearson correlation analysis was performed to analyze the relationship between *F. prausnitzii* number and butyrate. P-values and correlation coefficients are shown in the insert of each panel.

## References

1. Lopez-Siles, M., et al., Faecalibacterium prausnitzii: from microbiology to diagnostics and prognostics. ISME J, 2017. 11(4): p. 841–852.

2. Lopez-Siles, M., et al., Faecalibacterium prausnitzii: from microbiology to diagnostics and prognostics. The ISME Journal, 2017. 11(4): p. 841–852.

3. Zhao, H., et al., Systematic review and meta-analysis of the role of Faecalibacterium prausnitzii alteration in inflammatory bowel disease. Journal of Gastroenterology and Hepatology, 2021. 36(2): p. 320–328.

4. Furet, J.P., et al., Differential adaptation of human gut microbiota to bariatric surgery-induced weight loss: links with metabolic and low-grade inflammation markers. Diabetes, 2010. 59(12): p. 3049–57.

5. Karlsson, F.H., et al., Gut metagenome in European women with normal, impaired and diabetic glucose control. Nature, 2013. 498(7452): p. 99–103.

6. Qin, J., et al., A metagenome-wide association study of gut microbiota in type 2 diabetes. Nature, 2012. 490(7418): p. 55–60.

7. Zhang, X., et al., Human Gut Microbiota Changes Reveal the Progression of Glucose Intolerance. PLOS ONE, 2013. 8(8): p. e71108.

8. Da Silva, H.E., et al., Nonalcoholic fatty liver disease is associated with dysbiosis independent of body mass index and insulin resistance. Scientific Reports, 2018. 8(1): p. 1466.

9. Lopez-Siles, M., et al., Changes in the Abundance of Faecalibacterium prausnitzii Phylogroups I and II in the Intestinal Mucosa of Inflammatory Bowel Disease and Patients with Colorectal Cancer. Inflammatory Bowel Diseases, 2015. 22(1): p. 28–41.

10. Martín, R., L.G. Bermúdez-Humarán, and P. Langella, Searching for the Bacterial Effector: The Example of the Multi-Skilled Commensal Bacterium Faecalibacterium prausnitzii. Frontiers in Microbiology, 2018. 9(346).

11. Miquel, S., et al., Identification of Metabolic Signatures Linked to Anti-Inflammatory Effects of Faecalibacterium prausnitzii. mBio, 2015. 6(2): p. e00300–15.

12. Quévrain, E., et al., Identification of an anti-inflammatory protein from <em>Faecalibacterium prausnitzii</em>, a commensal bacterium deficient in Crohn’s disease. Gut, 2016. 65(3): p. 415–425.

13. Martín, R., et al., Faecalibacterium prausnitzii prevents physiological damages in a chronic low-grade inflammation murine model. BMC Microbiology, 2015. 15(1): p. 67.

14. Hill, C., et al., The International Scientific Association for Probiotics and Prebiotics consensus statement on the scope and appropriate use of the term probiotic. Nature Reviews Gastroenterology & Hepatology, 2014. 11(8): p. 506–514.

15. Palleja, A., et al., Recovery of gut microbiota of healthy adults following antibiotic exposure. Nature Microbiology, 2018. 3(11): p. 1255–1265.

16. David, L.A., et al., Diet rapidly and reproducibly alters the human gut microbiome. Nature, 2014. 505(7484): p. 559–63.

17. Mardinoglu, A., et al., An Integrated Understanding of the Rapid Metabolic Benefits of a Carbohydrate-Restricted Diet on Hepatic Steatosis in Humans. Cell Metabolism, 2018. 27(3): p. 559-571.e5.

18. Ruiz-Saavedra, S., et al., Comparison of Different Dietary Indices as Predictors of Inflammation, Oxidative Stress and Intestinal Microbiota in Middle-Aged and Elderly Subjects. Nutrients, 2020. 12(12).

19. Dewulf, E.M., et al., Insight into the prebiotic concept: lessons from an exploratory, double blind intervention study with inulin-type fructans in obese women. Gut, 2013. 62(8): p. 1112–1121.

20. Hustoft, T.N., et al., Effects of varying dietary content of fermentable short-chain carbohydrates on symptoms, fecal microenvironment, and cytokine profiles in patients with irritable bowel syndrome. Neurogastroenterology & Motility, 2017. 29(4): p. e12969.

21. Ramirez-Farias, C., et al., Effect of inulin on the human gut microbiota: stimulation of Bifidobacterium adolescentis and Faecalibacterium prausnitzii. British Journal of Nutrition, 2008. 101(4): p. 541–550.

22. Fernando, W., et al., Diets supplemented with chickpea or its main oligosaccharide component raffinose modify faecal microbial composition in healthy adults. Beneficial Microbes, 2010. 1(2): p. 197–207.

23. Hooda, S., et al., 454 Pyrosequencing Reveals a Shift in Fecal Microbiota of Healthy Adult Men Consuming Polydextrose or Soluble Corn Fiber. The Journal of Nutrition, 2012. 142(7): p. 1259–1265.

24. McDonald, D., et al., American Gut: an Open Platform for Citizen Science Microbiome Research. mSystems, 2018. 3(3): p. e00031–18.

25. Gonzalez, A., et al., Qiita: rapid, web-enabled microbiome meta-analysis. Nature Methods, 2018. 15(10): p. 796–798.

26. Amir, A., et al., Deblur Rapidly Resolves Single-Nucleotide Community Sequence Patterns. mSystems, 2017. 2(2): p. e00191–16.

27. Caporaso, J.G., et al., QIIME allows analysis of high-throughput community sequencing data. Nature Methods, 2010. 7(5): p. 335–336.

28. Duncan, S.H., et al., Growth requirements and fermentation products of Fusobacterium prausnitzii, and a proposal to reclassify it as Faecalibacterium prausnitzii gen. nov., comb. nov. Int J Syst Evol Microbiol, 2002. 52(Pt 6): p. 2141–2146.

29. Van den Abbeele, P., et al., Arabinoxylo-Oligosaccharides and Inulin Impact Inter-Individual Variation on Microbial Metabolism and Composition, Which Immunomodulates Human Cells. Journal of Agricultural and Food Chemistry, 2018. 66(5): p. 1121–1130.

30. Pérez-Burillo, S., et al., An in vitro batch fermentation protocol for studying the contribution of food to gut microbiota composition and functionality. Nature Protocols, 2021. 16(7): p. 3186–3209.

31. Nadkarni, M.A., et al., Determination of bacterial load by real-time PCR using a broad-range (universal) probe and primers set. Microbiology (Reading), 2002. 148(Pt 1): p. 257–266.

32. Lopez-Siles, M., et al., Mucosa-associated Faecalibacterium prausnitzii and Escherichia coli co-abundance can distinguish Irritable Bowel Syndrome and Inflammatory Bowel Disease phenotypes. Int J Med Microbiol, 2014. 304(3-4): p. 464–75.

33. Wishart, D.S., et al., HMDB 5.0: the Human Metabolome Database for 2022. Nucleic Acids Research, 2021. 50(D1): p. D622–D631.

34. Team, R.C., R: A language and environment for statistical computing. R Foundation for Statistical Computing. 2021.

35. Madrid-Gambin, F., et al., AlpsNMR: an R package for signal processing of fully untargeted NMR-based metabolomics. Bioinformatics, 2020. 36(9): p. 2943-2945.

36. !!! INVALID CITATION !!! [24].

37. Lundberg, S.M., et al., From local explanations to global understanding with explainable AI for trees. Nature Machine Intelligence, 2020. 2(1): p. 56–67.

38. Lopez-Siles, M., et al., Cultured Representatives of Two Major Phylogroups of Human Colonic Faecalibacterium prausnitzii Can Utilize Pectin, Uronic Acids, and Host-Derived Substrates for Growth. Applied and Environmental Microbiology, 2012. 78(2): p. 420–428.

39. Fitzgerald, C.B., et al., Comparative analysis of Faecalibacterium prausnitzii genomes shows a high level of genome plasticity and warrants separation into new species-level taxa. BMC Genomics, 2018. 19(1): p. 931.

40. Ruiz-Ojeda, F.J., et al., Effects of Sweeteners on the Gut Microbiota: A Review of Experimental Studies and Clinical Trials. Advances in Nutrition, 2019. 10(suppl_1): p. S31–S48.

41. Soto-Martin, E.C., et al., Vitamin Biosynthesis by Human Gut Butyrate-Producing Bacteria and Cross-Feeding in Synthetic Microbial Communities. mBio, 2020. 11(4): p. e00886–20.

42. Vital, M., A. Karch, and D.H. Pieper, Colonic Butyrate-Producing Communities in Humans: an Overview Using Omics Data. mSystems, 2017. 2(6): p. e00130–17.

43. De Filippis, F., E. Pasolli, and D. Ercolini, Newly Explored <em>Faecalibacterium</em> Diversity Is Connected to Age, Lifestyle, Geography, and Disease. Current Biology, 2020. 30(24): p. 4932-4943.e4.

44. Taylor, B.C., et al., Consumption of Fermented Foods Is Associated with Systematic Differences in the Gut Microbiome and Metabolome. mSystems, 2020. 5(2): p. e00901–19.

45. Cotillard, A., et al., A posteriori dietary patterns better explain variations of the gut microbiome than individual markers in the American Gut Project. The American Journal of Clinical Nutrition, 2021. 115(2): p. 432–443.

46. Cuesta-Zuluaga, J.d.l., et al., Age- and Sex-Dependent Patterns of Gut Microbial Diversity in Human Adults. mSystems, 2019. 4(4): p. e00261–19.

47. Zhu, C., et al., Determine independent gut microbiota-diseases association by eliminating the effects of human lifestyle factors. BMC Microbiology, 2022. 22(1): p. 4.

48. Moreno-Indias, I., et al., Red wine polyphenols modulate fecal microbiota and reduce markers of the metabolic syndrome in obese patients. Food & Function, 2016. 7(4): p. 1775–1787.

49. Clements, R.S., Jr and B. Darnell, Myo-inositol content of common foods: development of a high-myo-inositol diet. The American Journal of Clinical Nutrition, 1980. 33(9): p. 1954–1967.

50. Jiang, Z., et al., Dietary fruit and vegetable intake, gut microbiota, and type 2 diabetes: results from two large human cohort studies. BMC Medicine, 2020. 18(1): p. 371.

51. van Soest, A.P.M., et al., Associations between Pro- and Anti-Inflammatory Gastro-Intestinal Microbiota, Diet, and Cognitive Functioning in Dutch Healthy Older Adults: The NU-AGE Study. Nutrients, 2020. 12(11): p. 3471.

52. Sakamoto, M., et al., Genome-based, phenotypic and chemotaxonomic classification of Faecalibacterium strains: proposal of three novel species Faecalibacterium duncaniae sp. nov., Faecalibacterium hattorii sp. nov. and Faecalibacterium gallinarum sp. nov. International Journal of Systematic and Evolutionary Microbiology, 2022. 72(4).

53. Koecher, K.J., et al., Estimation and Interpretation of Fermentation in the Gut: Coupling Results from a 24 h Batch in Vitro System with Fecal Measurements from a Human Intervention Feeding Study Using Fructo-oligosaccharides, Inulin, Gum Acacia, and Pea Fiber. Journal of Agricultural and Food Chemistry, 2014. 62(6): p. 1332–1337.

54. Kim, H., et al., Co-Culture with Bifidobacterium catenulatum Improves the Growth, Gut Colonization, and Butyrate Production of Faecalibacterium prausnitzii: In Vitro and In Vivo Studies. Microorganisms, 2020. 8(5): p. 788.

55. Soto-Martin, E.C., et al., Vitamin Biosynthesis by Human Gut Butyrate-Producing Bacteria and Cross-Feeding in Synthetic Microbial Communities. mBio, 2020. 11(4): p. e00886–20.

56. Costabile, A., et al., A double-blind, placebo-controlled, cross-over study to establish the bifidogenic effect of a very-long-chain inulin extracted from globe artichoke (Cynara scolymus) in healthy human subjects. British Journal of Nutrition, 2010. 104(7): p. 1007–1017.

57. Ramnani, P., et al., Prebiotic effect of fruit and vegetable shots containing Jerusalem artichoke inulin: a human intervention study. British Journal of Nutrition, 2010. 104(2): p. 233–240.

