## Supplementary-Tables for "Application of computational data modeling to a large-scale population cohort assists the discovery of specific nutrients that influence beneficial human gut bacteria *Faecalibacterium prausnitzii*"

Table S1: Nutrients in the models

| Nutrient names in the models | Full name | unit |
|---|---|---|
| alcohol | Alcohol | g |
| inositol | Inositol | g |
| xylitol | Xylitol | g |
| sfa220 | SFA 22:0 (behenic acid) | g |
| delttoco | Delta-Tocopherol | mg |
| lycopene | Lycopene | mcg |
| alphacar | Alpha-Carotene (provitamin A carotenoid) | mcg |
| galactos | Galactose | g |
| vita_re | Total Vitamin A Activity (Retinol Equivalents) | mcg |
| sucrose | Sucrose | g |
| betaine | Betaine | mg |
| betacryp | Beta-Cryptoxanthin (provitamin A carotenoid) | mcg |
| vita_rae | Total Vitamin A Activity (Retinol Activity Equivalents) | mcg |
| thiamin | Thiamin (vitamin B1) | mg |
| coumest | Coumestrol | mg |
| maltose | Maltose | g |
| vitd3 | Vitamin D3 (cholecalciferol) | mcg |
| pfa183 | PUFA 18:3 (linolenic acid) | g |
| choline | Choline | mg |
| vita_iu | Total Vitamin A Activity (International Units) | IU |
| erythr | Erythritol | g |
| tfa182t | TRANS 18:2 (trans-octadecadienoic acid [linolelaidic acid]); includes c-t, t-c, t-t) | g |
| tagatose | Tagatose | mg |
| ribofla | Riboflavin (vitamin B2) | mg |
| addsugar | Added Sugars (by Available Carbohydrate) | g |
| vitd_iu | Vitamin D (calciferol) | IU |
| vitd | Vitamin D (calciferol) | mcg |
| pfa225 | PUFA 22:5 (docosapentaenoic acid [DPA]) | g |
| sorbitol | Sorbitol | g |
| adsugtot | Added Sugars (by Total Sugars) | g |

Table S2: Performance of various models

| Model ID | model_category | ml_algorithm | TARGET | Bin definition | target_label | train_roc_AUC | train_pr_AUC | train_number | test_roc_AUC | test_pr_AUC | test_number |
|---|---|---|---|---|---|---|---|---|---|---|---|
| A | low-high | RANDOM_FOREST | cube_trnfrmd_fprau | Q1 vs Q4 | low | 0.63 ± 0.01 | 0.66 ± 0.02 | 1532 | 0,61 | 0,65 | 386 |
| B | low-not_low | RANDOM_FOREST | cube_trnfrmd_fprau | Q1 vs Q2, Q3 and Q4 | low | 0.64 ± 0.01 | 0.41 ± 0.02 | 1554 | 0,64 | 0,41 | 764 |
| C | high-not_high | RANDOM_FOREST | cube_trnfrmd_fprau | Q1, Q2, and Q3 vs Q4 | high | 0.58 ± 0.02 | 0.3 ± 0.02 | 1532 | 0,61 | 0,32 | 764 |
| D | low-high | RANDOM_FOREST | cube_trnfrmd_fprau | < mean - 1SD vs > mean + 1SD | low | 0.66 ± 0.03 | 0.71 ± 0.03 | 828 | 0,64 | 0,71 | 216 |
| E | low-not_low | RANDOM_FOREST | cube_trnfrmd_fprau | < mean - 1SD vs rest | low | 0.65 ± 0.02 | 0.3 ± 0.03 | 896 | 0,68 | 0,33 | 764 |
| F | high-not_high | RANDOM_FOREST | cube_trnfrmd_fprau | > mean + 1SD vs rest | high | 0.56 ± 0.02 | 0.16 ± 0.02 | 828 | 0,58 | 0,17 | 764 |
| G | low-high | XGBOOST | cube_trnfrmd_fprau | < mean - 1SD vs > mean + 1SD | low | 0.65 ± 0.03 | 0.7 ± 0.03 | 828 | 0,65 | 0,73 | 216 |
| H | low-not_low | XGBOOST | cube_trnfrmd_fprau | < mean - 1SD vs rest | low | 0.66 ± 0.03 | 0.29 ± 0.03 | 896 | 0,65 | 0,32 | 764 |
| I | high-not_high | XGBOOST | cube_trnfrmd_fprau | > mean + 1SD vs rest | high | 0.56 ± 0.02 | 0.16 ± 0.02 | 828 | 0,58 | 0,17 | 764 |

Table S3: Summary of metadata of a subset of American Gut Project participants used in the study

| Summary of metadata of a subset of American Gut Project participants used in the study | |
|---|---|
| | **Overall (N=3816)** |
| **sex** | |
| female | 2268 (59.4%) |
| male | 1486 (38.9%) |
| other | 8 (0.2%) |
| unspecified | 54 (1.4%) |
| **age_cat** | |
| 20s | 261 (6.8%) |
| 30s | 520 (13.6%) |
| 40s | 725 (19.0%) |
| 50s | 833 (21.8%) |
| 60s | 1020 (26.7%) |
| 70+ | 253 (6.6%) |
| baby | 14 (0.4%) |
| child | 49 (1.3%) |
| teen | 36 (0.9%) |
| Unspecified | 105 (2.8%) |
| **host_age** | |
| Mean (SD) | 51.3 (15.6) |
| Median [Min, Max] | 53.0 [0.100, 173] |
| Missing | 104 (2.7%) |
| **host_height** | |
| Mean (SD) | 174 (67.3) |
| Median [Min, Max] | 170 [12.7, 1800] |
| Missing | 39 (1.0%) |
| **host_weight** | |
| Mean (SD) | 71.1 (17.9) |
| Median [Min, Max] | 69.0 [0.454, 214] |
| Missing | 37 (1.0%) |
| **host_body_mass_index** | |
| Mean (SD) | 31.3 (182) |

| | |
|---|---|
| Median [Min, Max] | 23.6 [0.200, 6890] |
| Missing | 53 (1.4%) |
| **bmi_cat** | |
| Normal | 2094 (54.9%) |
| Obese | 364 (9.5%) |
| Overweight | 1103 (28.9%) |
| Underweight | 167 (4.4%) |
| Unspecified | 88 (2.3%) |
| **race** | |
| African American | 20 (0.5%) |
| Asian or Pacific Islander | 108 (2.8%) |
| Caucasian | 3437 (90.1%) |
| Hispanic | 75 (2.0%) |
| Other | 106 (2.8%) |
| Unspecified | 70 (1.8%) |
| **country_residence** | |
| Australia | 56 (1.5%) |
| Austria | 4 (0.1%) |
| Belgium | 6 (0.2%) |
| Brazil | 1 (0.0%) |
| Canada | 33 (0.9%) |
| China | 2 (0.1%) |
| Czech Republic | 4 (0.1%) |
| Denmark | 5 (0.1%) |
| Estonia | 1 (0.0%) |
| France | 12 (0.3%) |
| Germany | 18 (0.5%) |
| Greece | 2 (0.1%) |
| Guernsey | 1 (0.0%) |
| Hong Kong | 1 (0.0%) |
| Ireland | 17 (0.4%) |
| Italy | 7 (0.2%) |
| Japan | 2 (0.1%) |
| Jersey | 2 (0.1%) |
| Mexico | 3 (0.1%) |
| Netherlands | 4 (0.1%) |
| New Zealand | 3 (0.1%) |
| Norway | 2 (0.1%) |
| Saudi Arabia | 1 (0.0%) |
| Serbia | 1 (0.0%) |
| Singapore | 1 (0.0%) |
| Slovakia | 4 (0.1%) |
| Spain | 6 (0.2%) |
| Sweden | 11 (0.3%) |
| Switzerland | 15 (0.4%) |
| United Arab Emirates | 4 (0.1%) |

| | |
|---|---|
| United Kingdom | 829 (21.7%) |
| United States | 1654 (43.3%) |
| United States Minor Outlying Islands | 1 (0.0%) |
| Unspecified | 1103 (28.9%) |
| **diet_type** | |
| Omnivore | 2929 (76.8%) |
| Omnivore but do not eat red meat | 247 (6.5%) |
| Unspecified | 97 (2.5%) |
| Vegan | 121 (3.2%) |
| Vegetarian | 185 (4.8%) |
| Vegetarian but eat seafood | 237 (6.2%) |
| **exercise_frequency** | |
| Daily | 811 (21.3%) |
| Never | 102 (2.7%) |
| Occasionally (1-2 times/week) | 1040 (27.3%) |
| Rarely (a few times/month) | 392 (10.3%) |
| Regularly (3-5 times/week) | 1397 (36.6%) |
| Unspecified | 74 (1.9%) |

Table S4: Mean intake of nutrients that are significantly different between the low and not low F. prausnitzii categories

Mean intake of nutrients that are significantly different between the low and not low F. prausnitzii categories

| nutrient | low (n=560) | not low (n=3256) | p.value | p.adj |
|---|---|---|---|---|
| alcohol | 9,31 | 13,72 | 0 | 0 |
| inositol | 1696,51 | 1944,21 | 0 | 0 |
| aspartam | 45,62 | 23,45 | 0 | 0 |
| betacryp | 10107,05 | 10987,59 | 0 | 0 |
| betacar | 159,3 | 175,29 | 0 | 0,01 |
| vita_iu | 0,14 | 0,16 | 0 | 0,01 |
| vita_re | 8240,81 | 8824,92 | 0 | 0,01 |
| alphacar | 4,28 | 4,56 | 0 | 0,02 |
| pectins | 19797,68 | 21525,04 | 0 | 0,02 |
| vita_rae | 1341,38 | 1437,73 | 0 | 0,02 |
| lutzeax | 2260,96 | 2441,68 | 0 | 0,04 |

Wilcoxon rank sum test was used to compare the two groups and Benjamini-Hochsberg method was applied to control the false discovery rate. Only significant are shown (adjusted $p<0.05$).

Table S5: Normalized mean intake (2000 kcal) of nutrients that are significantly different between the low and not low F. prausnitzii categories

| Normalized mean intake (2000 kcal) of nutrients that are significantly different between the low and not low F. prausnitzii categories | | | | |
|---|---|---|---|---|
| nutrient | low (n=560) | not low (n=3256) | p.value | p.adj |
| alcohol | 9,87 | 14,75 | 0 | 0 |
| inositol | 2040,82 | 2284,09 | 0 | 0,0000028 |
| proline | 48,44 | 25,9 | 0,0000064 | 0,000309 |
| aspartam | 186,62 | 203,74 | 0,0000105 | 0,0003791 |
| glutamic | 2,37 | 2,24 | 0,000045 | 0,0013057 |
| betacryp | 11,1 | 10,6 | 0,0002905 | 0,0070194 |
| delttoco | 15,77 | 14,98 | 0,0005321 | 0,0110212 |
| gammtoco | 0,16 | 0,18 | 0,0018005 | 0,0290081 |
| thiamin | 5,13 | 4,8 | 0,001611 | 0,0290081 |
| selenium | 119,48 | 114,55 | 0,0020092 | 0,0291329 |
| alphacar | 1,54 | 1,46 | 0,0026063 | 0,0343557 |

Wilcoxon rank sum test was used to compare the two groups and Benjamini-Hochsberg method was applied to control the false discovery rate. Only significant are shown (adjusted $p<0.05$).
